# Redesign of energetically frustrated regions rescues function in defective T4 clamp loaders

**DOI:** 10.64898/2026.05.08.723874

**Authors:** Siddharth Nimkar, Thu Nguyen, Deepti Karandur, Subu Subramanian, Michael E O’Donnell, John Kuriyan

## Abstract

DNA polymerase clamp loaders are AAA+ ATPases that load sliding clamps on DNA for high-speed replication. Using a platform for high-throughput mutagenesis of replication proteins in T4 bacteriophage, we carried out saturation mutagenesis of the AAA+ ATPase module of the T4 clamp loader bearing a mutation, Gln 118◊Asn (Q118N), that reduces fitness. We identified residues for which different mutations improve the fitness of the Q118N variant but are neutral in the wild-type background. These conditionally neutral “rescue hotspots” overlap with those identified earlier in another defective variant (D110C). These rescue hotspots localize to regions where the sequence is not optimal for the structure, as determined by energetic frustration analysis. We designed new sequences for three of these regions, using the protein-design algorithm ProteinMPNN. In two helical regions, several designed sequences increased the fitness of both wild-type and mutant proteins, likely due to enhanced stability. An inter-domain hinge in AAA+ module changes conformation during activation, and designs for the hinge lead to loss of fitness in the wild-type background. However, when using the active conformation as the template, designs for the hinge increase the fitness of defective variants. In contrast designs templated on the inactive conformation led to loss of fitness, suggesting that a proper conformational balance is crucial. Thus, adaptive capacity in the clamp loader resides in a network of conditionally neutral sites that enable functional tuning through shifts in stability and conformational equilibria.

## Introduction

The versatility of the ATP-driven molecular machines that are essential for cellular function is remarkable (Mulkidjanian et al. 2007). The specialization and diversification of these machines is made possible by their capacity to tolerate mutations, enabling adaptation without catastrophic loss of function. DNA polymerase clamp loaders, which are oligomeric ATPases responsible for loading sliding DNA clamps onto DNA for high-speed processive replication, are a useful system for addressing these questions (Kelch et al. 2012). Clamp-loader activity depends on integrating ATP binding, DNA recognition, and sliding clamp opening (Kelch et al. 2012). Clamp loaders are members of the very large AAA+ family of oligomeric ATPases, whose members play critical roles in diverse aspects of cellular mechanism (Neuwald et al. 1999; Khan et al. 2022). Studying the adaptive capacity of clamp loaders is therefore of broad interest in terms of understanding how the complexity and functional specialization of an important class of molecular machines arose.

We have developed a robust high-throughput mutagenesis and in-vitro evolution platform for measuring the fitness of the replication proteins of T4 bacteriophage in their natural context (Subramanian et al. 2021; Subramanian et al. 2024). The platform takes advantage of the fact that T4 phage utilize their own DNA replication proteins, encoded in the phage genome, including a replicative DNA polymerase and a clamp-loader/clamp system (Benkovic and Spiering 2017). We have used this platform previously to map the mutational fitness landscape of the T4 clamp loader and the sliding clamp (Subramanian et al. 2021; Marcus et al. 2024; Subramanian et al. 2024).

Clamp-loader complexes across all domains of life are pentameric ATPase assemblies (Jeruzalmi et al. 2002; Kelch et al. 2012; Carver et al. 2025). In T4 phage, the clamp-loader complex contains four copies of the gp44 ATPase subunit, encoded by gene 44, and a fifth subunit, denoted gp62, that has lost the ATPase domain. The five positions in the clamp loader complex are labeled A, B, C, D, and E (Kelch et al. 2012). The four ATPase subunits are at the B, C, D, and E positions in the assembly. The ATPase subunits consist of N-terminal AAA+ modules, comprised of a RecA-type (Story and Steitz 1992). ATPase Domain 1 and a helical Domain 2 that is characteristic of AAA+ proteins (Guenther et al. 1997; Lenzen et al. 1998; Yu et al. 1998). Gp62 is at the A position, and it is referred to as the “clasp” subunit because its V-shaped structure bridges the subunits at B and E (Figure 1A). Each of the five subunits of the complex has a C-terminal collar domain, and these oligomerize to form the pentameric assembly. ATP binding and hydrolysis power a cycle in which the clamp loader recognizes and opens the sliding clamp and loads it on primer-template DNA (Guenther et al. 1997; Jeruzalmi et al. 2001; Bowman et al. 2004; Kelch et al. 2011; Marzahn et al. 2014; Zheng et al. 2022).

**Fig. 1.**
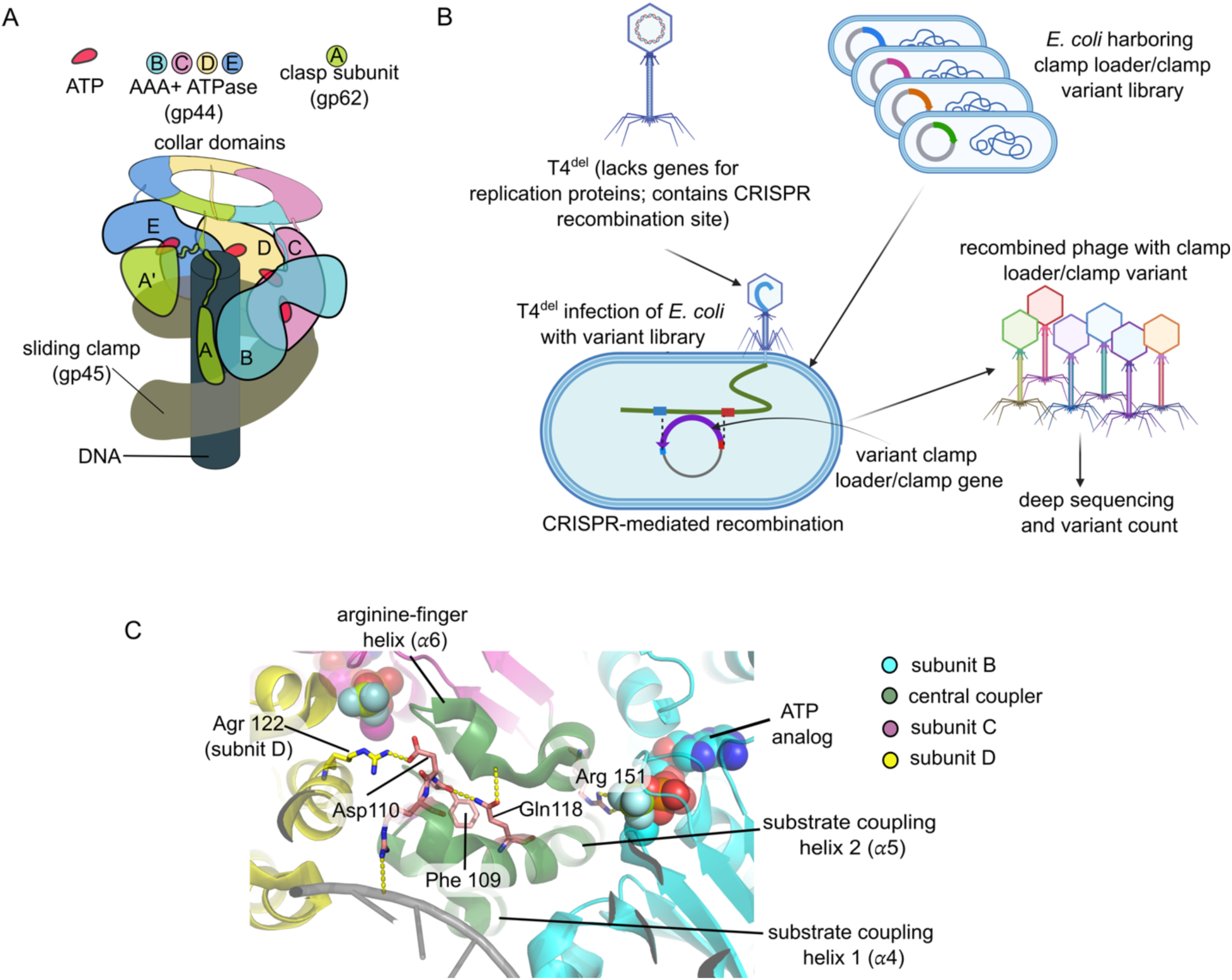
The T4 clamp loader complex and the phage propagation assay. a) Schematic diagram of the T4 clamp loader (gp44 and gp62) bound to DNA and the sliding clamp. b) The phage-propagation assay. The T4^del^ variant of bacteriophage T4 lacks the genes for the clamp loader and the sliding clamp (genes 44, 62, and 45). A plasmid library containing clamp-loader variants is transformed into *E. coli* prior to infection with T4^del^ phage. The clamp loader and clamp genes are inserted into the phage genome by CRISPR-mediated recombination, enabling phage replication. The resulting phage are harvested, and the fitness of each variant is determined through deep sequencing (Subramanian et al. 2021). c) Close-up view of the clamp loader structure showing the locations of Gln 110 and Asp 110, respectively

Our high-throughput assay for clamp-loader function (Subramanian et al. 2021) utilizes a modified T4 phage, referred to as T4^del^, in which the genes encoding the sliding clamp (gene 45) and the clamp-loader subunits (genes 44 and 62) have been deleted (Figure 1B). T4^del^ phage are competent to infect *Escherichia coli* cells, but the absence of clamp loader and clamp genes prevents phage replication. For high-throughput mutagenesis, we create a library of *E. coli* cells in which each cell carries a plasmid encoding a different variant of the clamp loader and clamp genes. Upon infection by T4^del^ these genes are inserted into the phage genome using a CRISPR-Cas12a system, thereby “repairing” the phage. Lysis of the *E. coli* cells then produces new phage particles whose replicative fitness in subsequent rounds of infection depends on the fitness of the clamp loader and clamp genes that were inserted into the phage during the first infection.

We determine the fitness of each variant by using Illumina sequencing to measure the abundance of each variant gene before phage infection and after cycles of infection are complete. The reinsertion of the replication genes into the phage genome by the CRISPR/Cas12a system occurs at the endogenous locus, and so fitness values measured in these experiments are obtained in the natural context, with natural levels of protein expression.

Despite the essential nature of the clamp-loading function, we found that the majority of residues in the clamp-loader complex and the sliding clamp are remarkably tolerant to substitution, with only a limited set of positions in the catalytic core and interaction interfaces being strongly constrained (Subramanian et al. 2021). Although in vitro deep mutagenesis experiments can mask the true sensitivity of sites to mutation (Rockah-Shmuel et al. 2015), for the T4 clamp loader/clamp system the mutational tolerance is highly correlated with lack of sequence conservation across different phages, indicating that the tolerance reflects an intrinsic property of the proteins (Subramanian et al. 2021).

The evolutionary implications of this mutational tolerance were highlighted in our subsequent studies of clamp-loader variants with diminished function. In one case, we engineered a severely defective chimeric clamp loader by replacing the AAA+ module of the T4 clamp loader with that from a divergent phage, *Aeromonas salmonicida* phage 44RR2.8t (denoted RR2 phage) (Nolan et al. 2006; Petrov et al. 2006; Subramanian et al. 2024). The fitness of modified T4 phage bearing this chimeric T4/RR2 clamp loader is ∼5000-fold lower than that of wild-type T4 phage (Subramanian et al. 2024). The T4/RR2 chimeric clamp loader recovers substantial fitness through single mutations that enhance interactions with DNA or the sliding clamp. Notably, many of these compensatory mutations occur at sites where mutations are close to neutral in the wild-type background, revealing that the clamp loader has a capacity to accumulate cryptic or latent variation that can potentially confer fitness advantages under perturbed conditions (Subramanian et al. 2024). This phenomenon highlights the role of cryptic variation and conditional neutrality in enabling rapid functional adaptation (Wagner 2005; Draghi et al. 2010; Hayden et al. 2011; Raman et al. 2016).

A complementary picture emerges from analysis of a mildly defective clamp loader bearing a mutation in the conserved DExD-box motif of the AAA+ module (Marcus et al. 2024). The last aspartate residue in this motif, Asp 110, forms an interfacial ion pair between adjacent subunits, and is important for coupling DNA binding to ATP hydrolysis (Figure 1C) (Kelch et al. 2011; Mallam et al. 2012). Replacement of Asp 110 by cysteine (D110C) reduces the fitness of the clamp-loader by ∼5-fold. Saturation mutagenesis of this variant identified “rescue hotspots” distributed across the AAA+ module, which are sites at which different substitutions restore function to varying degrees, despite having little or no effect on fitness of the wild-type protein. Cryo-EM analyses showed that in the absence of DNA, the T4 clamp loader adopts an autoinhibited conformation in which DNA binding is blocked and the interfacial catalytic sites are disrupted (Marcus et al. 2024). Some of the rescue hotspots are in regions that undergo conformational changes during activation, suggesting that they act by shifting the balance between inactive and active states.

These findings showed that the clamp loader harbors a distributed capacity for adaptation, rooted in sites that are not strongly constrained under normal conditions but become functionally important when the system is perturbed. However, key questions remain unresolved. What structural features distinguish such sites from those that are strictly constrained? How are they organized within the protein, and what physical principles govern their ability to modulate function? More broadly, can these latent adaptive features be identified systematically, and can they be leveraged to engineer improved function?

Here, we address these questions by analyzing the mutational landscape of the T4 clamp loader in another sensitized genetic background. Specifically, we mutated a highly conserved glutamine residue in the AAA+ module (Gln 118) that plays a key role in stabilizing the structural elements that couple ATP binding to DNA and clamp recognition (Figure 1C)(Subramanian et al. 2021; Sosa et al. 2026). Substitution of Gln 118 by asparagine (Q118N) reduces fitness by ∼20-fold (Subramanian et al. 2021). We now show that saturation mutagenesis in the context of the Q118N mutation reveals rescue hotspots that overlap with those identified in the D110C study.

We find that the rescue hotspots tend to be found in regions of high “energetic frustration” (Ferreiro et al. 2011; Treuheit et al. 2011; Fuglestad et al. 2013; Parra and Ferreiro 2025). In such regions, the energy of the native structure is not well separated from the energies of alternative structures, indicating that these regions are poised to undergo conformational change or accommodate sequence variation. We used the machine learning–based protein design program ProteinMPNN (Dauparas et al. 2022; Wicky et al. 2022) to generate new sequences for short segments in the hot-spot regions, based on the structures of active or inactive clamp loaders. For two regions, which are helical, we identified designed sequences that improved the fitness of the clamp loader beyond wild-type levels, most likely by stabilizing the structure. The third region is the inter-domain hinge. In this area, only designs based on the active conformation improve fitness, as mutations in the hotspot regions appear to alter the conformational balance of the clamp loader.

## Results and Discussion

### Saturation-mutagenesis screen for the AAA+ module in the Q118N background

To generate fitness data for double mutations within the AAA+ module of the T4 clamp loader (residues 2–230), we modified a previously developed single-site saturation mutagenesis library (Subramanian et al. 2021). Specifically, we introduced the Q118N mutation across this existing library to serve as a fixed genetic background. We then screened this modified library for effects on phage growth using the established T4 phage propagation assay (Subramanian et al. 2021; Marcus et al. 2024; Subramanian et al. 2024) . This approach allowed us to assess the fitness consequences of replacing each residue in the AAA+ module with all 20 amino acids and a stop codon, one at a time, in the presence of the Q118N mutation.

The fitness of each variant was measured by using Illumina sequencing to determine the abundance of each variant in the input library (input count) and in the phage population after bacterial infection and lysis (output count), as done previously (Subramanian et al. 2021). The fitness of a variant is defined as:

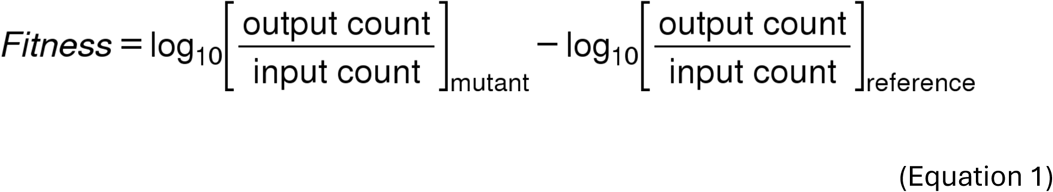

Here, mutant refers to a particular mutation for which the fitness is being measured, in the background of the fixed Q118N mutation. The reference sequence corresponds to phage with only the Q118N mutation in the AAA+ module. The experiment was repeated three times, and the fitness values were averaged over these replicates. A fitness value of +1 corresponds to 10-fold faster rate of propagation of that variant with respect to the Q118N mutant, for which the fitness value is zero. Note that the Q118N mutation has a fitness of - 1.25 when the wild-type clamp loader is used as the reference, corresponding to about a 20- fold reduction in growth rate with respect to wild-type phage (Figure S1B) (Subramanian et al. 2021)

As for the D110C variant (Marcus et al. 2024), the Q118N mutant clamp loader is more sensitive to mutations than the wild-type clamp loader, with most substitutions showing a decrease in fitness (Figure 2). The Q118N and D110C mutations are expected to destabilize the clamp loader because they disrupt key interactions. The incorporation of additional destabilizing mutations, such as replacement of residues in the hydrophobic core, will increase the probability that the clamp loader fails to fold and assemble. This decreased margin of stability potentiates the effects of destabilizing mutations elsewhere (Bloom et al. 2007). Nevertheless, the saturation mutagenesis data identify several mutations that increase the fitness of the Q118N variant (Figure 2 & S1C).

**Fig. 2.**
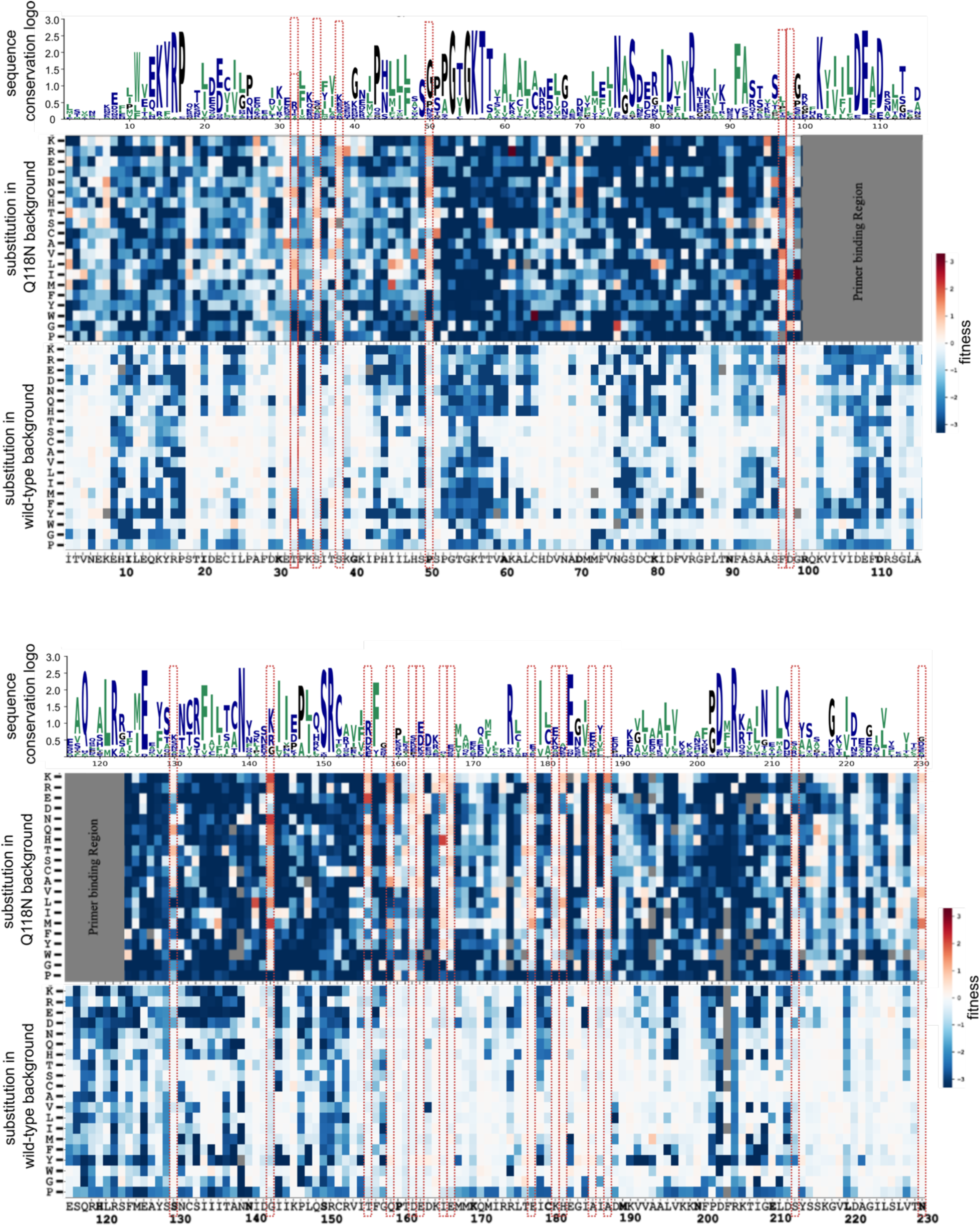
Saturation mutagenesis of Q118N T4 clamp loader. Shown here are heatmaps corresponding to fitness values obtained for saturation mutagenesis of the AAA+ module (residues 2 to 230 of gp44) of the Q118N mutant T4 clamp loader (top panel of each diagram). For comparison, saturation mutagenesis data for the wild-type T4 clamp loader are also shown, remeasured as part of this work (bottom panel of each diagram). The fitness values are indicated by a color scale, ranging from loss-of-fitness (blue) to neutral (white) to gain-of-fitness (red). The sequence conservation at each position is shown above the heatmaps, based on an alignment of ∼900 phage gp44 sequences (see Methods). The height of the letters indicates the frequency with which the corresponding amino acid is found at that position in the alignment. Red vertical boxes indicate the locations of "rescue hotspots"—positions where more than five amino acid substitutions improve fitness in the Q118N background. The grey box denotes an unmutated primer- binding region.

We define gain-of-function or “rescue mutations” as those that have fitness values >0.2, corresponding to a ∼1.67 fold or greater increase in phage growth over the Q118N mutant. At several sites where rescue mutations occur, more than one kind of mutation results in an increase in fitness – a similar result was obtained previously for saturation mutagenesis in the background of the D110C mutation (Marcus et al. 2024). We refer to sites that have more than five such rescue mutations as “rescue hotspots”.

### Rescue hotspot mutations are conditionally neutral

There are 14 rescue hotspots that are common to the D110C and Q118N saturation-mutagenesis datasets (Figure 3A and 3B). Mutations at these sites are neutral, or near neutral, in the wild-type background. The capacity of these mutations to increase fitness is revealed only upon the introduction of weakening mutations elsewhere, such as Q118N or D110C. The rescue hotspot mutations are therefore examples of *conditionally neutral mutations* that have little or no effect in the context of the wild-type protein, but can impact fitness when operating conditions are altered (Masel 2006; Bloom et al. 2007; Soskine and Tawfik 2010; Raman et al. 2016; Jayaraman et al. 2022).

**Fig. 3.**
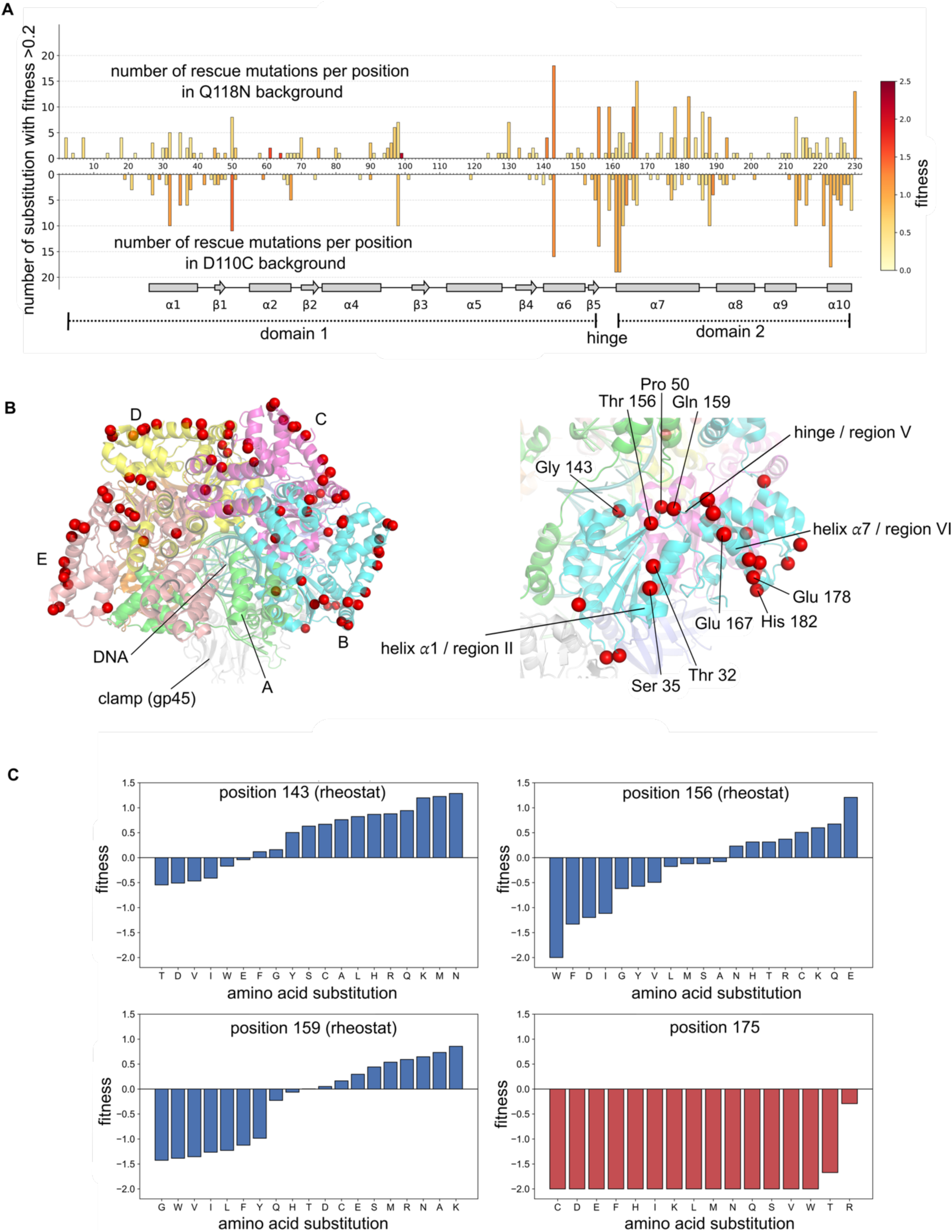
Location of rescue hotspots in the Q118N and D110C backgrounds. a) The graph shows the number of different amino acid substitutions that rescue function (fitness > 0.2) in the Q118N background at each position in the AAA+ module. For each rescue hotspot, the maximum value of the fitness for substitutions at that site are shown by a color gradient, with darker red representing increasing fitness. Rescue mutations in the D110C background, measured previously (Marcus et al. 2024), are shown in the diagram below. For ease of comparison, the vertical scale is flipped for the D110C data. The secondary structural elements of the AAA+ module are indicated below the diagram. b) View of the active T4 clamp-loader complex bound to primer-template DNA (PDB ID: 8uh7). The five subunits are denoted A through E, and the collar domains are not shown. Rescue hotspots that are common to the Q118N and D110C backgrounds are shown as red spheres. A detailed view focusing on subunit B is shown in the right. c) Bar plots detailing the fitness effects of all single amino acid substitutions at representative rescue hotspot positions (residues 143, 156 and 159) within Q118N clamp loader. This variable, substitution-dependent response defines these sites as "rheostat positions”(Meinhardt et al. 2013; Fenton et al. 2020). Position 175 is one of the positions that represents a highly sensitive residue where most of the amino acid substitution leads to loss of fitness (also refer Figure S2).

The rescue hotspots occur at sites of low sequence conservation, as shown by an alignment of ∼900 phage clamp-loader sequences (Figure 2). For example, Pro 50, one of the most prominent hotspots in both the Q118N and D110C datasets, is frequently replaced by asparagine, alanine, or aspartate in other phage clamp loaders. Saturation mutagenesis in the wild-type background shows that all substitutions at this site result in little or no change in fitness (Subramanian et al. 2021) (Figure 2).

Different substitutions at the rescue hotspots have different effects on fitness in the Q118N or D110C backgrounds, ranging from loss of fitness to gain of fitness (Figure 3C and S2). Analysis of mutational data for other proteins led to the concept of “rheostat positions” in proteins, where different substitutions result in a range of fitness effects (Meinhardt et al. 2013; Fenton et al. 2020; Swint-Kruse et al. 2021). Rheostat positions are not conserved – in contrast to strongly conserved sites that usually result in substantial loss of function when mutated (Figure 3C). The rescue hotspots are *conditional rheostat positions*, in that their capacity to serve as rheostats is only revealed when a detrimental mutation is introduced elsewhere.

Most of the rescue hotspot sites have no obvious roles in function or structural integrity, which is consistent with the features of rheostat positions in other proteins (Fenton et al. 2020). One exception is Gly 143, a prominent rescue hotspot in both the Q118N and D110C datasets, for which rationalization of gain-of-function mutations is straightforward. Most phage clamp loaders have an arginine or a lysine at this position, instead of glycine. This site is located near DNA, and arginine or lysine residues at this position could readily engage the phosphate backbone of DNA, thereby increasing DNA affinity (Marcus et al. 2024). In the Q118N and D110C backgrounds, rescue mutations at this position indeed include arginine and lysine. Glycine is also important for maintaining an inactive conformation of the T4 clamp loader (Marcus et al. 2024). Substitutions at this position, not just to arginine or lysine, are expected to lead to steric clash in the inactive conformation, thereby activating the clamp loader by destabilizing the inactive structure.

Gly 143 mutations are neutral, rather than gain-of-function, in the wild-type background (Subramanian et al. 2021). This result is consistent with results obtained previously using a clamp-loader variant with a severe defect in function. This variant has the AAA+ module of the T4 clamp loader replaced by that of another phage, RR2 (Subramanian et al. 2024). Saturation mutagenesis of the clasp subunit and the sliding clamp, which are retained from T4 in the T4/RR2 chimera, showed previously that many mutations that are neutral or near neutral in the wild-type background are gain-of-function in the chimeric clamp-loader background. These sites are conditional rheostat positions, where mutations increase fitness to varying degrees, presumably through varying effects on the affinity of the clamp loader for DNA or the sliding clamp (Subramanian et al. 2024).

### Rescue hotspots occur at sites of energetic frustration in the AAA+ module

The fact that different substitutions at the rescue hotspots result in increased fitness suggests that the wild-type residues at these sites are not optimal when deleterious mutations are introduced elsewhere. We turned to protein frustration analysis to gain insight into this phenomenon (Ferreiro et al. 2007; Ferreiro et al. 2011). In the energy landscape theory of protein folding, the principle of minimal frustration posits that the energy of the native structure of a protein is sufficiently well separated from the energies of alternative structures so that folding proceeds rapidly down an energetic funnel (Bryngelson et al. 1995). While the overall structure of a stable protein will be minimally frustrated, local regions can have high frustration – in these regions, the sequence of the protein does not define the local structure with sufficient energetic separation from alternative conformations. Such sequences allow the protein to undergo conformational changes, transmit allosteric signals, interact with binding partners, or accommodate mutations (Ferreiro et al. 2007; Ferreiro et al. 2011; Ferreiro et al. 2018).

Local frustration is defined by calculating the interaction energies of pairs of residues in the folded structure of the protein and comparing these energies to the energies of decoy pairs, sampled from the inter-residue distance and sequence probability distributions corresponding to the protein (Ferreiro et al. 2007). If the energy of an interacting pair is not well separated from the energies of interacting pairs in the decoys, the interaction is considered to be frustrated. A per-residue “configurational frustration index” is calculated based on all the pairwise interactions made by that residue within a cutoff distance, and the calculation is repeated for each residue to obtain a residue-dependent frustration index. The frustration index for a residue reflects the density of frustrated interactions made by that residue (Ferreiro et al. 2007; Parra et al. 2016; Rausch et al. 2021)

We used a web-based tool, the Frustratometer (Parra et al. 2016; Rausch et al. 2021), to calculate the local frustration indices for the AAA+ module of the T4 clamp loader. For this calculation we used the structure of the clamp-loader complex in the active state, bound to the sliding clamp, primer-template DNA, and an ATP analog (PDB 8UH7; (Kelch et al. 2011;

Marcus et al. 2024)). Although each of the three ATPase subunits of the clamp loader have a different disposition to DNA and the sliding clamp, they exhibit similar patterns of local frustration (Figure S3). There are seven regions of high local frustration, labeled regions I to VII in Figure 4.

**Fig. 4.**
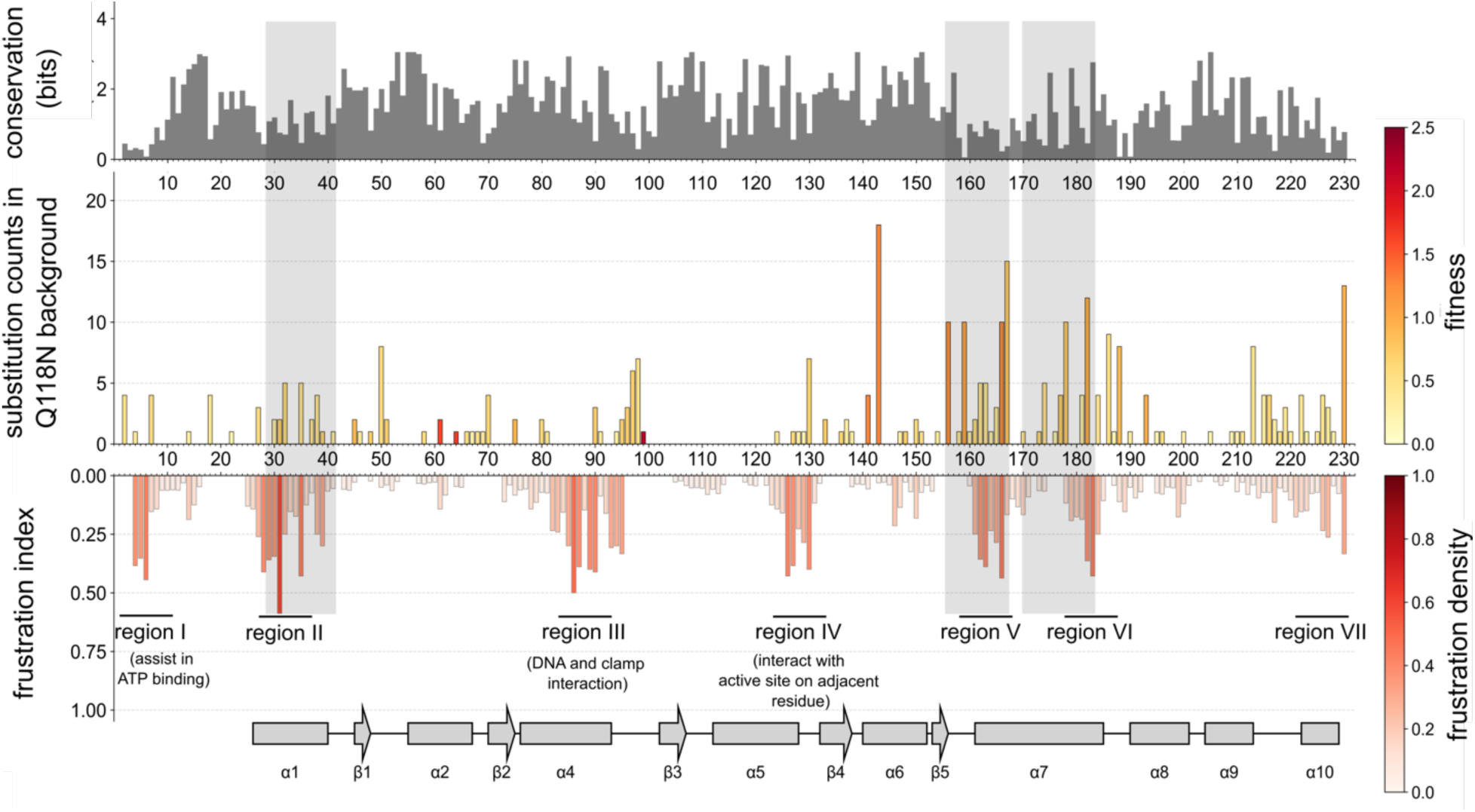
Local energetic frustration and rescue hotspot locations. The diagram shows the distribution of rescue hotspots in the Q118N background, as in Figure 3. Energetic frustration index, calculated for Subunit B of the active clamp loader complex (PDB ID: 8uh7). The downward-facing bars represent the density of highly frustrated interactions within a 5 Å radius of the reference residue, with warmer colors indicating higher frustration density. Calculated using the Frustratometer (Parra et al. 2016; Rausch et al. 2021), and using the configurational frustration index as defined previously. The sequence conservation of the AAA+ module and the secondary structural elements are also shown. The grey shading indicates regions chosen for protein sequence design by ProteinMPNN (refer Figure 5). Rescue hotspots occur in regions of low sequence conservation. For helix 𝛼7, the amphipathicity of the helix places highly conserved residues facing the hydrophobic core. The rescue hotspots are on the hydrophilic and less conserved face of the helix.

Rescue hotspots are sparse in frustrated regions I, III, and IV, which are highly conserved in sequence and have important functional roles. Region I (residues 5 to 15) is important for linking ATP binding to inter-subunit interactions in the clamp loader. Region III (residues 76 to 95) is part of the central coupler, a core element of the allosteric machinery that couples DNA binding to clamp recognition (Subramanian et al. 2021). Region IV (residues 121-130) is also part of the central coupler, and it links ATP binding to DNA recognition. Gln 118, the critical residue that is mutated in the Q118N saturation-mutagenesis experiments, is adjacent to this region. We infer that although regions I, III, and IV exhibit high local frustration, they do not readily accommodate mutations because of their important functional roles.

Rescue hotspots tend to occur in frustrated regions II, V, VI, and VII. Region V (residues 159 to 170) spans the hinge connecting domains 1 and 2 of the AAA+ module. Upon activation, the AAA+ module undergoes a hinge-bending motion that alters the conformation of the hinge (region V). This conformational change converts the ATPase active site from catalytically inactive to active (Marcus et al. 2024). Region V is not conserved in sequence and is tolerant of mutation in the wild-type context (Figure 4). We assume that the rescue mutations in this region alter the conformational bias of the hinge, thereby compensating for the deleterious effects of the Q118N or D110C mutations.

The involvement of frustrated regions II, VI, and VII in the functional cycle is more subtle. Like region V, they are not conserved in sequence and are mutationally tolerant in the wild-type context. All three regions span a helices, and so do not undergo significant internal changes during clamp-loader function. All three frustrated regions do, however, change their interactions with neighboring domains or subunits during the inactive to active transition, and so mutations in these regions could alter the balance between these states. Region II (residues 27-40; helix 𝛼1) is located on the outer surface of the clamp loader, near the AAA+ module of an adjacent subunit, and the loop preceding this helix recognizes the adenine ring of ATP or ADP. Region VI (residues178-185) corresponds to the C-terminal portion of helix 𝛼7, which transduces changes in the hinge (region V) into movement of Domain 2 of the AAA+ module with respect to Domain 1. Region VII (helix 𝛼10) is the last helix of the AAA+ module, and it connects to the collar domain via a 5-residue linker. In the structures of inactive states of the T4 clamp loader the AAA+ modules of two of the subunits cannot be visualized, suggesting that the transition from active (DNA bound) to inactive (DNA free) releases strain in the organization of the AAA+ modules with respect to the collar domain, which will affect Region VII.

### Using the protein design program ProteinMPNN to alter sequences around rescue hotspots

We used the protein design program ProteinMPNN (Dauparas et al. 2022) to investigate whether computational sequence design can be used to increase the fitness of wild type and mutant clamp loaders (Figure 5A). Given a structural template, ProteinMPNN generates sequences that are compatible with that template. In contrast to earlier design methods that used energy calculations to select optimal sequences, ProteinMPNN relies on machine-learning algorithms that match sequences to local structure based on patterns learnt from known protein structures. ProteinMPNN has demonstrated an impressive ability to generate sequences that result in well folded proteins and has been applied successfully to design completely new protein folds (Dauparas et al. 2022; Wicky et al. 2022).

**Fig. 5.**
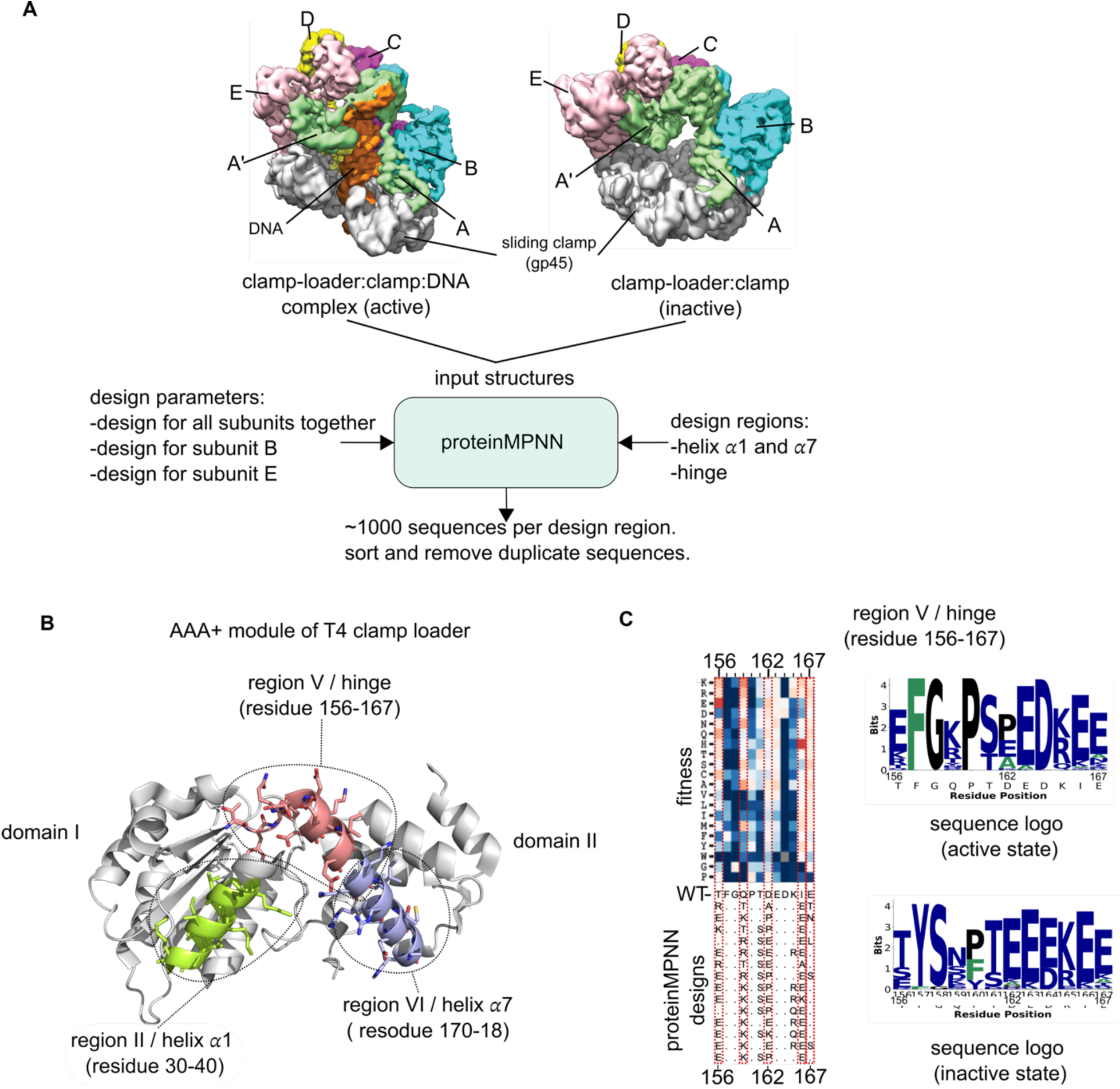
Using ProteinMPNN to design segments of the AAA+ module. a) The workflow for the ProteinMPNN-based design process. The structures of the DNA-bound (active, PDB: 8uh7) and DNA-free (inactive, PDB: 8unh) T4 clamp loader were utilized as input templates. Approximately 900 sequences were generated per region and filtered to isolate unique sequences. b) Structure of the B subunit from the active clamp loader complex (PDB: 8uh7), showing the three frustrated regions for which designs were generated —region II, (helix 𝛼1; residues 30- 40), region V (the inter-domain hinge; residues 156-167), and region VI (helix 𝛼7; residues 170-182). c) Alignment of ProteinMPNN designs with experimental mutational data for the Q118N mutant. The heatmap displays fitness data from saturation mutagenesis of the Q118N mutant. Below the heatmap, an alignment of 15 representative sequences designed using the active structure template is shown. Red dashed boxes highlight the rescue hotspots. Sequence logos corresponding to the distribution of amino acids in the designed sequences are also shown, for designs using the active (top) and inactive (bottom) structural templates.

We used the structures of active and inactive forms of the T4 clamp loader as templates for ProteinMPNN, and generated sequences for residues 30 to 40 in frustrated region II (helix 𝛼1), residues 156 to 167 in region V (the hinge), and residues 170 to 182 in region VI (C-terminal end of helix 𝛼7) (Figure 5B). We did not model region VII. The number of unique sequences generated by ProteinMPNN depends on the ‘temperature’ of the design calculation, with higher temperatures yielding more designed sequences, but with decreased reliability (Dauparas et al. 2022). For the active template, we obtained 256, 450, and 350 unique sequences for the three regions, respectively. For the inactive template, we obtained 297, 457, and 558 sequences for the three regions. A sampling of the results of the ProteinMPNN calculations with inactive and active templates is shown in Figure 5C and S4. For all three frustrated regions, the positions at which ProteinMPNN changes the sequence tend to be rescue hotspots in the Q118N and D110C saturation-mutagenesis experiments.

We made DNA libraries of clamp-loader variants in which the gene encoding the ATPase subunit (gene 44) had the segments corresponding to frustrated regions II, V, or VI replaced by codons corresponding to the ProteinMPNN-designed sequences, with only one segment replaced in any one gene. Separate libraries were constructed for the wild-type clamp loader as well as for the Q118N and D110C mutants. Each library contained designs based on the active and the inactive templates. Plasmids encoding the library were transfected into *E. coli* cells that were then infected with the T4^del^ phage that lacks clamp-loader and clamp genes. The relative fitness of each design was determined using the phage-propagation assay. Each experiment was repeated three or more times, and the fitness scores were averaged over replicates.

For experiments involving protein designs, each sequence was synthesized as part of an oligonucleotide pool (Twist Biosciences Inc), and the wild-type or reference DNA is present at levels close to that of each variant. Due to the stochastic nature of phage infection, we observed occasional “jackpot events” (Luria and Delbrück 1943), characterized by disproportionate representation of certain sequences in the output of the experiment. These jackpot events arise due to stochastic timing differences in the production of the first set of progeny phage, which are then exponentially amplified. This phenomenon can lead to a less fit phage taking over the population due an accident of early infection. Because the jackpot events are random, averaging of fitness values across multiple independent runs mitigates against this effect. For the fitness distributions considered below, the standard deviation of fitness values in each range of fitness provides a guide to reliability. Jackpot events may be mitigated by spiking in an excess of the wild-type clamp loader DNA.

### Effect of ProteinMPNN designs for the hinge region of the AAA+ module

The principal conformational change undergone by the T4 clamp loader as it activates is a rearrangement of the AAA+ modules with respect to each other and to the collar and clasp domains (Marcus et al. 2024). In addition, within each AAA+ module there is a change in the orientation of domain 1 with respect to domain 2, which is accompanied by a change in the conformation of the interdomain hinge.

We created three libraries with ProteinMPNN designs for the interdomain hinge (residues 156 to 160, which spans the hinge and first 7 residues in helix 𝛼7) (Figure 6A). In one, the ProteinMPNN-designed segments were embedded in the wild-type clamp loader sequence (the wild-type hinge library). In the other two, the designs were embedded in the Q118N and D110C clamp loader sequences (the Q118N and D110C hinge libraries). The reference sequence included in the wild-type library is the wild-type sequence, and for the other two libraries the Q118N and D110C variants of the clamp loader were included as reference sequences, respectively.

**Fig. 6.**
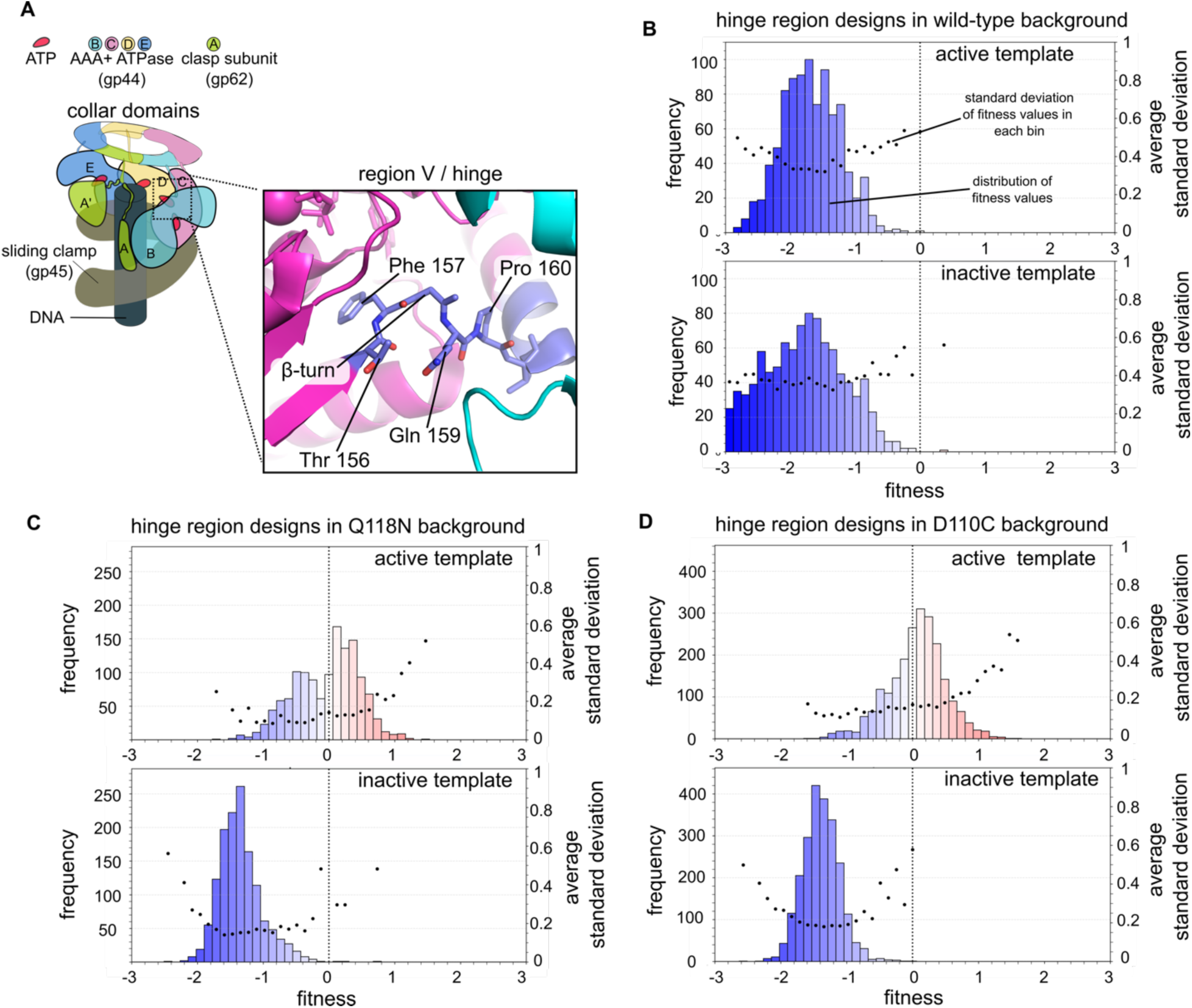
Effects on fitness of ProteinMPNN designs for the hinge. a) Structure of the inter-domain hinge (residues156-160). b) Distribution of fitness scores for ProteinMPNN-designed hinge sequences in the wild-type clamp loader background. The average standard deviation for the measurements in each fitness bin is also shown. c) and d) distribution of fitness scores for hinge designs introduced into the Q118N and D110C mutant backgrounds, respectively.

Figure 6B shows distributions of fitness obtained for the wild-type hinge libraries for designs with the active and inactive templates. The fitness distributions are shown as frequency histograms, with each bar representing the frequency with which phage variants with a particular fitness value are observed in the experiment. The frequency is calculated by aggregating the results of three independent runs of the phage propagation assay, and the standard deviations for the fitness values in each bin are also shown.

For the wild-type clamp loader, replacement of the hinge region by the ProteinMPNN designs results in substantial loss-of-function, for designs with both active and inactive templates (Figure 6B). The distributions are peaked at fitness values near -1.5 for both active and inactive designs, corresponding to a ∼30-fold reduction in phage propagation rates with respect to the wild-type clamp loader. There are no designs with significant gain of fitness over wild type.

For the Q118N hinge library, most designs with the inactive template led to substantial loss of fitness, with the distribution again peaked at about -1.5, corresponding to a 30-fold decrease in phage propagation rate with respect to the reference Q118N phage (Figure 6C). For results with designs based on the active template, the distribution of fitness effects is centered around zero (no loss of fitness), and about 55% of the designs using the active template result in gain-of-fitness with respect to the Q118N reference. About 12% of the designs result in fitness values greater than 0.5, corresponding to a 3-fold increase in phage propagation rate with reference to the Q118N mutant. Similar results were obtained with the D110C hinge library (Figure 6D).

### Stabilization of the hinge in the active conformation can recover fitness in clamp loaders with deleterious mutations

The ability to undergo conformational change is critical for function, and the balance between the active and inactive states is, presumably, tuned appropriately in the wild-type clamp loader. We infer that optimizing the hinge sequence for either the active or inactive states, as done in the wild-type hinge library, results in loss of fitness as these designs cause the equilibrium to become unbalanced, leading to a reduction in the ability of the clamp loader to cycle between states. In the case of the Q118N and D110C clamp loaders we assume that the initial mutations cause the equilibria to be displaced towards the inactive state. Introduction of hinge designs with the inactive templates into these backgrounds further shift the equilibrium to the inactive state, causing more substantial loss-of-function. In contrast, we assume that hinge designs based on the active template shift the equilibrium towards the active state, correcting the imbalance.

To further explore this idea, we turned to the chimeric RR2/T4 clamp loader, in which the AAA+ module is replaced by that of the clamp loader from RR2 phage (Figure 7A). The AAA+ module of the RR2 clamp loader is 64% identical to that of T4 phage. Phage with the chimeric clamp loader exhibit a 5000-fold reduction in propagation rate with respect to wild-type T4 phage (Subramanian et al. 2024). The introduction of a single mutation (D86H) in the RR2 AAA+ module decreases electrostatic repulsion between the chimeric clamp loader and the T4 sliding clamp, and the phage propagation rate improves so that it is only 20-fold lower than for wild-type T4 phage (Subramanian et al. 2024).

**Fig. 7.**
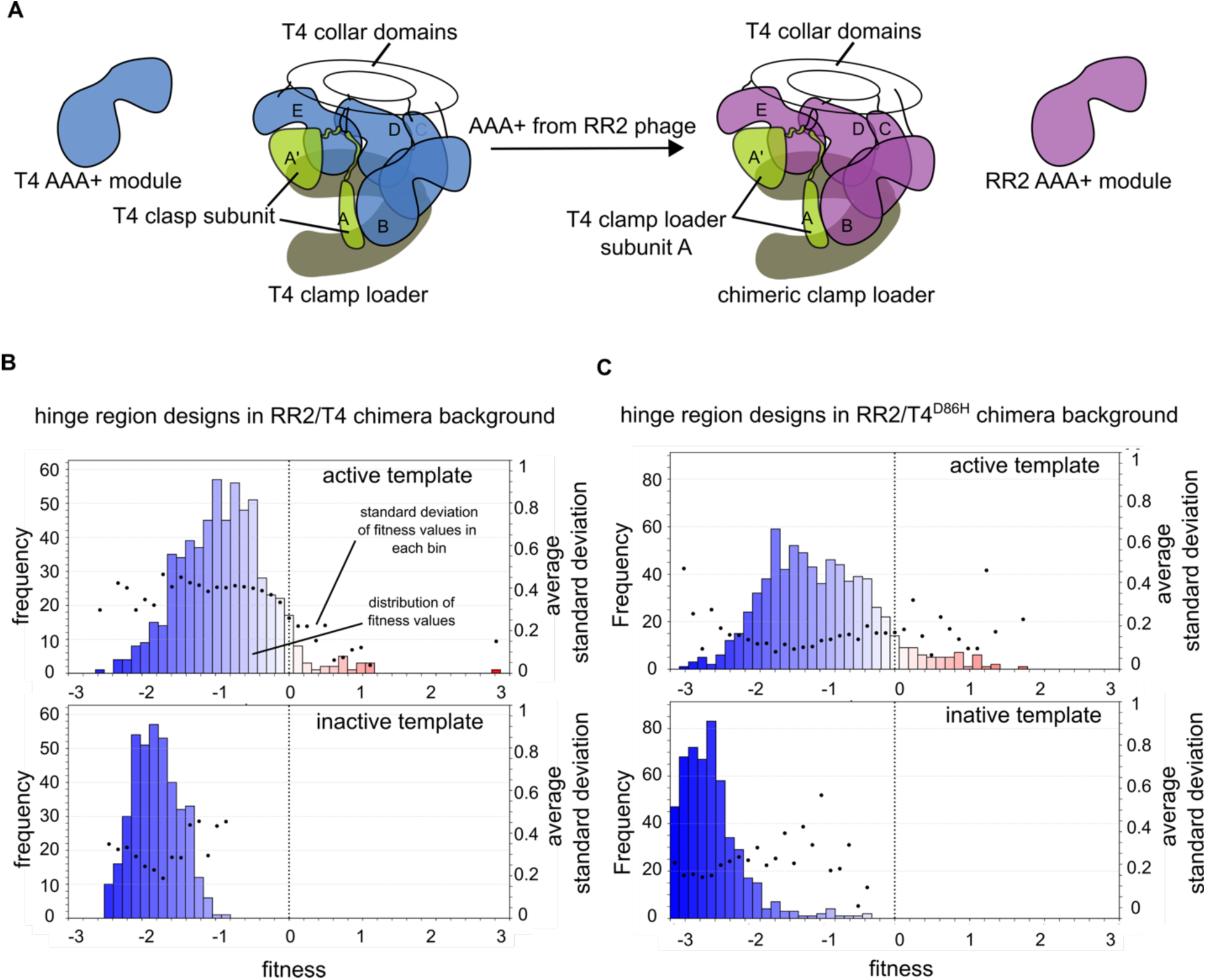
Effect of ProteinMPNN designs for the hinge in the T4/RR2 chimeric clamp loader. a) Schematic representation of the T4/RR2 chimeric clamp loader. The AAA+ modules of subunits B–E from the T4 clamp loader (blue) are replaced by the AAA+ modules from the RR2 clamp loader (purple). The T4 clasp subunit A (green) and collar domains are retained from the T4 clamp loader, as is the sliding clamp (gp45). b) Fitness distributions for ProteinMPNN-designed hinge sequences embedded in the T4/RR2 chimera. The hinge designs were templated on the structure of the wild-type T4 clamp loader and are the same as those used in the experiments shown in Figure 6. c) Fitness distributions for hinge designs embedded in the T4/RR2^D86H^ chimera, a mutant with partially restored fitness (Subramanian et al. 2024).

Data from saturation mutagenesis of the T4/RR2 chimera, done previously (Subramanian et al. 2024) and the T4/RR2^D86H^ variant (this work; Figure S5) show that there are fewer rescue mutations than seen for the Q118N or D110C mutant clamp loaders. This increased sensitivity to mutation is likely due to reduced stability of the T4/RR2 chimeric clamp loader, as suggested by studies on other proteins (Hidalgo et al. 2022). Attempts to purify the T4/RR2 and D86H clamp-loader complexes yielded very little protein, consistent with reduced stability of the chimeric protein complexes.

We investigated whether multi-residue designs generated by ProteinMPNN could improve the fitness of the T4/RR2 chimeras. We generated two libraries (the T4/RR2 hinge library and the T4/RR2 ^D86H^ hinge library) using the same ProteinMPNN designs used for experiments with the T4 clamp loader. We carried out phage competition experiments with these two libraries, using phage with the T4/RR2 and the T4/RR2^D86H^ chimeric clamp loaders as the reference sequences, respectively, and with two replicates each.

For both sets of experiments, ProteinMPNN designs templated on the inactive structure led to substantial loss of fitness (∼100-fold to ∼1000-fold reduction in phage propagation rates with respect to T4/RR2 or T4/RR2^D86H^). When ProteinMPNN designs templated on the active structure are used instead, the fitness distribution improves, with distributions for both T4/RR2 and T4/RR2^D86H^ centered around fitness values of -1 (a ten-fold reduction in fitness with respect to the reference). In both backgrounds, some of the ProteinMPNN designs show significant gain-of-fitness with respect to the reference sequences (Figure 7B & C). This supports the idea that ProteinMPNN designs templated on the active structure can shift the conformational equilibrium of AAA+ module to the active state.

To understand the difference between the ProteinMPNN designs templated on the active and inactive structures we looked at the distributions of amino acids at each position in the hinge for designs for the T4 clamp loader bearing the Q118N mutation (Figure S6). The most obvious characteristic of the designs that increase fitness is the incorporation of an ion pair between residues 156 and 159 (Figure 6A). The residues at these positions in the wild-type T4 clamp loader are threonine and glutamine, respectively, and the sidechains point towards each other, but do not appear to interact. These residues are at positions *i* and *i*+3 of a type II b turn, and the geometry of the b turn sets up the sidechains at these positions for potential interaction (Wilmot and Thornton 1988). All the designs that increase fitness have a glutamate at position 156 and a lysine or an arginine at position 159. These residues are well positioned to form an ion pair that would stabilize the b turn.

A type II b turn strongly favors a glycine residue at the *i*+2 position (residue 158), due to steric clash with the carbonyl group of the *i*+1 residue (Wilmot and Thornton 1988). The wild-type T4 clamp loader has a glycine at this position, as does the RR2 clamp loader, and all of the designs that work maintain a glycine at this position, emphasizing the importance of the b turn. The b turn positions Phe 157 so that the sidechain is very tightly packed in the hydrophobic core, and all the designs that work preserve the identity of this sidechain as well. The designs also preserve proline at position 160, a residue that is important for maintaining a sharp bend that connects the b turn to the first helix (𝛼7) of domain 2 (Figure S6).

The designs that do not work have features that would be expected to destabilize the b turn or the tight connection to helix 𝛼7 (Figure S6). Many of the sequences lack the ion pair between positions *i* and *i*+3 of the b turn, and some do not preserve the glycine residue at position *i*+2. Phe 157 is replaced by leucine or tyrosine in 60% of the sequences, and Pro 160 by phenylalanine or tyrosine in 40% of the sequences (Figure S6). The conformational change that occurs on transitioning from the active to the inactive conformation loosens up the packing of residues around the b turn, and the ProteinMPNN designs appear to reflect this fact.

We also examined ProteinMPNN-designed sequences that increased fitness for the T4/RR2 and T4/RR2^D86H^ chimeras (Figure S7). The features exhibited by these sequences are very similar to those seen for sequences that rescue the T4^Q118N^ clamp loader, including the introduction of an ion pair that stabilizes the b turn. The RR2 clamp loader has alanine at position 160, and the sequences that improve fitness replace this residue by proline, as seen in the T4 clamp loader. These results further support the hypothesis that stabilizing the active conformation of the clamp loader can improve the fitness of defective clamp loader variants.

### ProteinMPNN designs for helices 𝛼1 and 𝛼7 can improve the fitness of the wild-type clamp loader

Fitness values were measured for the T4 clamp loader with frustrated region II (helix 𝛼1) replaced with ProteinMPNN-designed sequences, and Figure 8A shows histograms for the fitness scores obtained for the wild-type libraries for designs with the active and inactive templates. The graphs also show the standard deviations of the fitness measurements in each bin of data. Considering designs obtained using the active template, a striking result is that ∼18% of the designs result in fitness values that are improved with respect to the wild-type clamp loader (Figure 8A). The gain-of-function in the wild-type background was unexpected because the single-site saturation mutagenesis data for the clamp loader did not reveal any gain-of-function mutations in the wild-type background (Subramanian et al. 2021). In contrast to the saturation mutagenesis experiments, where mutations are introduced one at a time, the ProteinMPNN designs implemented here replace up to 11 residues at a time.

**Fig. 8.**
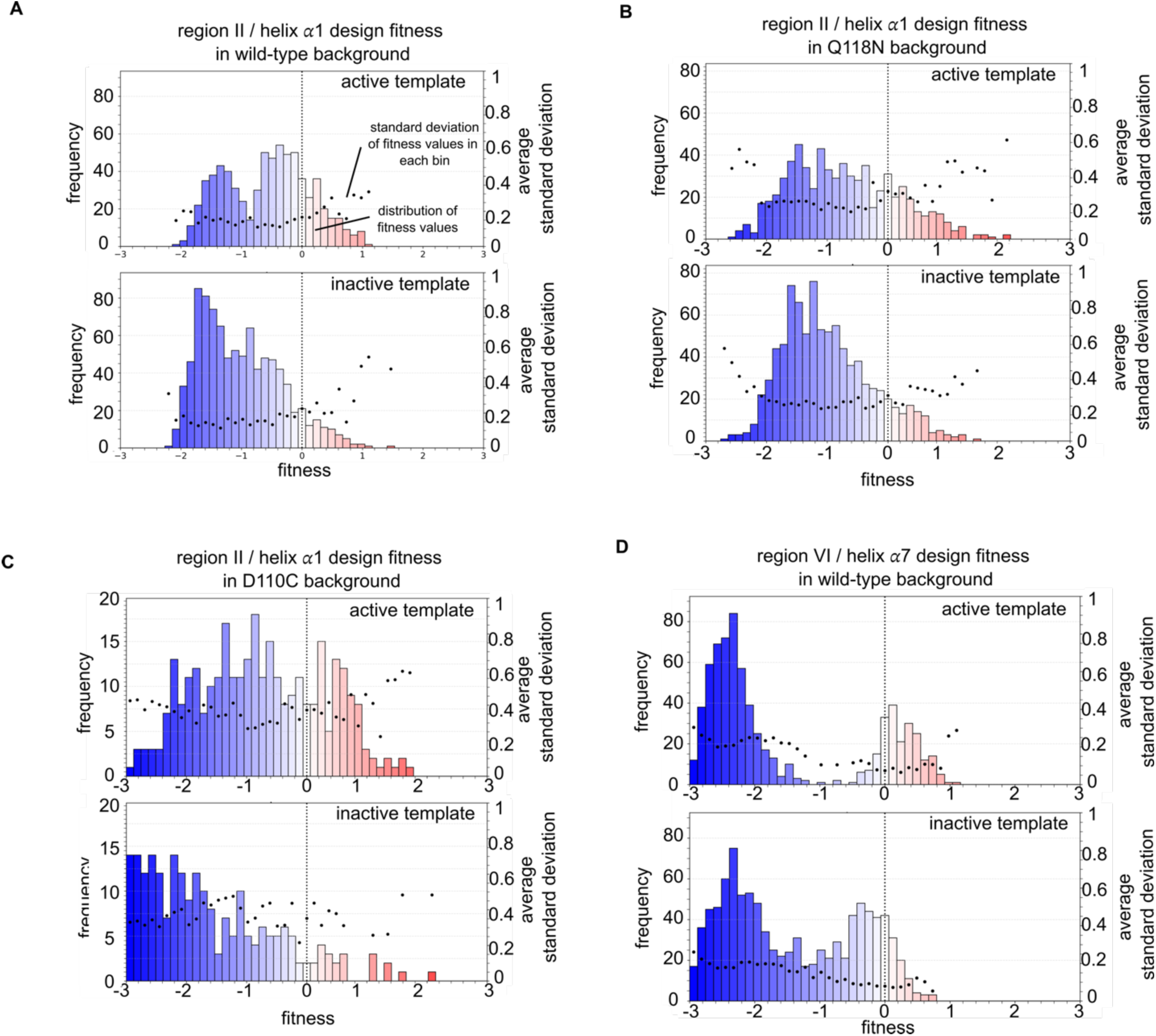
ProteinMPNN designs for region II (helix 𝛼1) and region VI (C-terminal portion of helix 𝛼7) Distribution of fitness scores for ProteinMPNN designs for region II (helix 𝛼1) embedded in a) the wild-type background, b) the Q118N background, and c) the D110C background. d) Distribution of fitness scores for region VI (C-terminal portion of helix 𝛼7), embedded in the wild-type background.

The distributions of fitness values for designs using the inactive templates resembles the distribution obtained using the active template. This is consistent with the fact that there is no significant internal structural change in this region between the inactive and active states, and so the templates for the design are similar in terms of local structure. The distribution does shift to lower fitness values when the inactive template is used, with a lower fraction of variants (∼8% compared to 18%) displaying fitness values greater than for the wild-type clamp loader.

We also measured fitness effects for ProteinMPNN designs for frustrated region II (helix 𝛼1) in the background of the Q118N mutation and D110C mutations and obtained qualitatively similar results (Figure 8B & C). The reference sequences for these two libraries are the Q118N and D110C variants of the clamp loader, respectively. For Q118N, ∼22% and ∼12% of the designs using the active template and inactive templates, respectively, led to improvements in fitness with respect to Q118N mutant. For D110C, ∼16% and ∼5% of the designs templated on the active and inactive structures led to improvements in fitness with respect to the D110C mutant. Taken together, these results suggest that the fitness effects have two components – some of the increase in fitness is likely due to increasing the stability of the clamp loader complex, and some might be due to preferential stabilization of the active conformation.

We also created a library of ProteinMPNN designs for frustrated region VI (helix 𝛼7) in the wild-type T4 clamp loader background (Figure 8D). The results using this library are similar to those obtained for region II. For designs using the active template, ∼13% of the sequences result in fitness increases over wild type. For designs using the inactive template, the distribution is shifted towards lower fitness, with ∼4% of the sequences resulting in improvements over wild-type fitness.

To understand why the ProteinMPNN sequences result in increased fitness we compared the helical propensities of the wild-type and the designed sequences. For Region II (helix 𝛼1), the ProteinMPNN designs substitute surface residues in the wild-type helix with alternates that have higher helical propensity (Deléage and Roux 1987) (Figure S8). This includes replacement of threonine by leucine or valine, serine by glutamate or alanine, and lysine by arginine. For region VI (helix 𝛼7) there is a general trend towards increased helical propensity, but there are also some correlated changes involving tertiary interactions. For example, in one design with increased fitness, Glu 178 is replaced by leucine, which reduces helical propensity but increases hydrophobic interaction with Ile 2, located directly below Glu 178. A correlated change is the replacement of Lys 181, which forms an ion pair with Glu 178 in the wild-type structure, by glutamate, which increases the helical propensity and also sets up an ion pair with His 182.

Taken together, these results suggest that the fitness effects arising from ProteinMPNN designs for helix 𝛼1 and 𝛼7 are due to increasing the stability of the clamp loader complex. However, the shift towards increased fitness values in the histograms for the designs based on the active templates suggest that there might also be some preferential stabilization of the active conformation.

## Conclusions

In this work we have analyzed the mutational response of the AAA+ module of the T4 clamp loader to two mutations that weaken the stability of the active conformation of the complex. The first mutation, D110C, replaces an aspartate residue in the conserved DExD box and thereby weakens the ability of the clamp loader to form the fully configured ATP-bound complex, and has been studied previously (Marcus et al. 2024). In the present work we have carried out saturation mutagenesis of the Q118N mutant form of the clamp loader. The replacement of Gln 118 by asparagine compromises a network of hydrogen bonds that stabilizes the central coupler, a unit in the AAA+ module that couples ATP binding to DNA recognition and the sliding clamp (Subramanian et al. 2021; Sosa et al. 2026).

A key result is that saturation mutagenesis of the Q118N variant identifies a number of rescue hotspots in the AAA+ module that are common to those identified previously for the D110C variant. These hotspots are characterized by their tolerance to mutation in the wild-type background and by the presence of multiple substitutions capable of improving fitness when a deleterious mutation is introduced elsewhere, indicating that they function as rheostat-like positions (Fenton et al. 2020). Importantly, their effects are conditional: they remain largely silent in the context of the wild-type protein but become functionally significant when the system is perturbed. Our findings support a model where the T4 clamp loader possesses a latent adaptive capacity, whereby mutations that are silent in the wild-type context reveal the ability to restore structure stability or alter conformational balance when a destabilizing mutation is introduced elsewhere (Figure 9A).

**Fig. 9.**
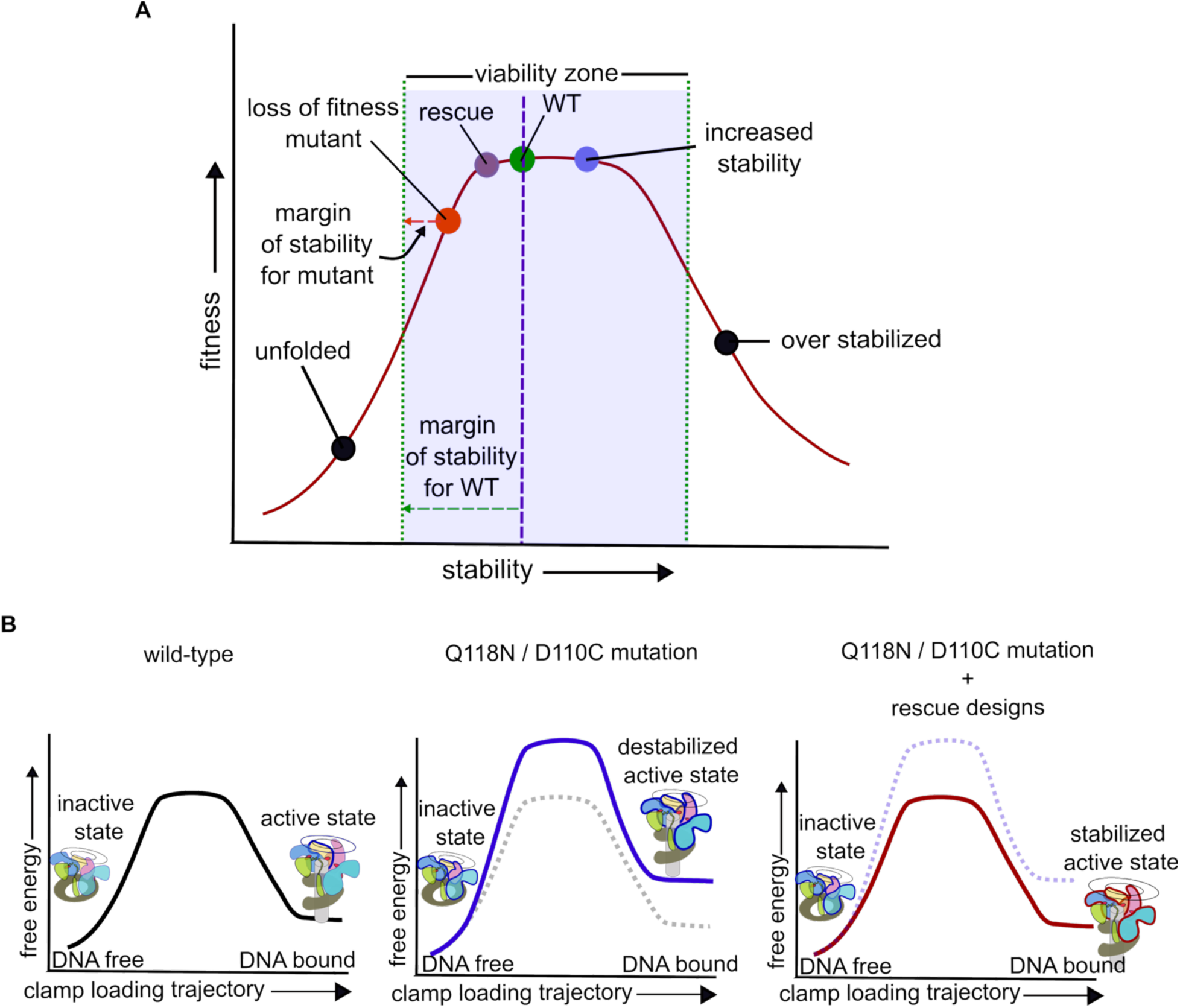
Latent adaptive capacity mitigates fitness loss by restoring the margin of stability. a) The relationship between structural stability and functional fitness is governed by a critical "margin of stability" required for optimal activity. The wild-type (WT) protein operates at peak fitness safely within this thermodynamic threshold. When deleterious mutations disrupt this balance, the protein shifts outside the stability margin, leading to a loss of activity (red). Compensatory sequence design can rescue these defective variants (purple) by restoring stability. Conversely, introducing stabilizing designs into the already-stable wild-type background (blue) shifts the complex further to the right. This could result in improved fitness but could also result in loss of fitness due to conformational rigidity. b) Conceptual free-energy landscape for the clamp loading cycle is depicted. Protein sequence designs that favor the active state (red line) rescue the function of the destabilized clamp loader (adapted from Figure 6e in (Marcus et al. 2024) ).

Analysis of these rescue hotspots shows that they occur in regions identified as energetically frustrated. Frustrated regions appear to encode a degree of sequence and structural ambiguity, allowing changes in these regions to modulate conformational equilibria or inter-domain interactions (Ferreiro et al. 2011). While some highly frustrated regions (such as those involved in core allosteric coupling) are functionally constrained and do not tolerate mutation, others—particularly regions II (helix 𝛼1), V (the hinge), VI (the C-terminal portion of helix 𝛼7), and VII (helix 𝛼10)—are both frustrated and permissive, making them sites where mutations can tune function.

We used ProteinMPNN (Dauparas et al. 2022) to design new sequences for three of the regions of high local frustration and found that some of these designed sequences can improve the fitness of the clamp loader. In frustrated regions II (helix 𝛼1) and VI (helix 𝛼7), multi-residue designs improve fitness not only in the Q118N and D110C backgrounds but also in the wild-type clamp loader. To understand the origin of these improvements, we must distinguish between two contributions to biological fitness: the "margin of stability" (the extent to which the protein structure is stabilized over the unfolding threshold) and the conformational equilibrium between inactive and active structural states. Our data indicate that while designs in the hinge region rescue fitness by stabilizing the active conformation (Figure 9B), the designs in regions II and VI likely improve fitness by also increasing the margin of stability.

Designs targeting the hinge region (region V) underscore the importance of maintaining a finely tuned conformational balance in the AAA+ module. In the wild-type background, designs templated on both the active- and inactive-states are deleterious, suggesting that the natural sequence for this region is optimized to support a balanced conformational equilibrium. However, in the Q118N and D110C backgrounds, designs based on the active conformation can restore fitness (Figure 9B). In contrast, designs based on the inactive conformation fail to improve fitness, suggesting that these designs shift the equilibrium toward inactive states. This interpretation is further reinforced by analysis of the successful designs, which reveal specific sequence features—most notably the introduction of an ion pair within a type II β-turn—that stabilize the conformation of the hinge in the active state.

Finally, experiments with the severely compromised T4/RR2 chimeric clamp loader demonstrate that these principles extend to more extreme perturbations. Although this chimera exhibits reduced stability and a diminished capacity for rescue through point mutations, ProteinMPNN designs targeting the hinge region can still yield measurable improvements in fitness, when based on the active conformation. The recurrence of similar sequence features across rescued variants in different genetic backgrounds emphasizes the generality of the underlying structural mechanisms.

Our results support a view of protein evolution in which adaptability is encoded in a distributed network of conditionally neutral sites that are energetically frustrated. Such sites enable proteins to respond to perturbations through modulation of conformational landscapes and adjustment of stability. Our work illustrates how latent adaptive capacities in a protein machine can be identified by saturation mutagenesis, offering avenues for understanding and engineering complex molecular machines. The fact that the rescue mutations do not have a discernable phenotype in the wildtype background indicates that the stability of the wild-type protein is more than sufficient for proper function, and it enables accommodation of a range of mutations that only show phenotypic signatures in a weakened background. The extent to which evolution has selected for this mutational robustness and responsive capacity merits further study.

## Materials and Methods

### Saturation Mutagenesis of the Q118N mutant T4 Clamp-Loader

For saturation mutagenesis of the AAA+ module of the ATPase subunit (gp44), we used a previously generated wild-type saturation mutagenesis library, in which every residue in the AAA+ module was replaced by each of the 20 amino acids, plus the stop codon, one at a time (Subramanian et al. 2021; Marcus et al. 2024). The library was split into two pools: in the first pool (pool A), residues 2–118 were varied, and in the second pool (pool B) residues 116–230 were varied.

The Q118N mutation was introduced on top of the existing mutations using primer-guided mutagenesis. Primers spanning from 20 nucleotides downstream of position 118 to 10 nucleotides upstream were used to introduce a mutagenetic codon for asparagine at position 118. The 5′ ends of these primers contained a recognition motif for the BsaI restriction enzyme for Golden Gate cloning (Engler et al. 2008). As a result the residues flanking the mutation site were not varied, and retained the wild-type identity. Library diversity during the cloning stages was maintained at >100X coverage per variant and was assessed by colony counts of transformants. The two library pools were used to transform *E. coli* (BL21) and used in the phage-propagation assay.

Parallel saturation mutagenesis of the wild-type gp44 AAA+ module was performed as an internal control. The resulting fitness landscape exhibited high concordance with previously published deep mutational scanning data (Subramanian et al. 2021).

### Phage-propagation assay

The phage-propagation assay was carried out as described previously (Subramanian et al. 2021). Plasmid libraries containing the engineered clamp-loader variants were electroporated into *E. coli* BL21 cells, achieving a transformation coverage exceeding 200- fold per variant. These host cells harbored a CRISPR-Cas12a plasmid specifically directing cleavage at a predefined *cas12a* recognition site engineered into the genome of the T4^del^ phage (Subramanian et al. 2021). To initiate exponential growth, roughly 50 µL of a saturated overnight culture was used to inoculate approximately 100 mL of fresh Luria-Bertani (LB) medium.

Once the culture reached an optical density (OD600) of 0.1, a 5 ml aliquot was withdrawn for plasmid extraction; DNA isolated from this sample provided the input counts for the calculation of fitness. The remaining culture was split into three flasks of 25 ml each for replicate experiments. T4^del^ phage, in which the genes for the clamp loader subunits (genes 44 and 62) and the sliding clamp (gene 45) are deleted, came from stocks prepared for earlier experiments (Subramanian et al. 2021). T4^del^ phage was introduced at a multiplicity of infection (MOI) of 0.001 (approximately 10⁸ phage particles for 10^11^ *E. coli* cells) to trigger infection and propagation. The infected cells were incubated at 37°C with continuous agitation for roughly 18 hours. The flasks were then left undisturbed at room temperature to allow for the gravity-driven sedimentation of cellular debris. A 1 ml sample of the culture was subsequently centrifuged to completely pellet any remaining debris, and the clarified supernatant was collected for extraction of DNA for determining the output counts for each variant.

To prepare the recombinant libraries for Illumina sequencing while minimizing potential PCR-induced biases, we utilized a three-stage nested PCR protocol restricted to 20 cycles per stage. Initially, the 3.6 kb recombinant locus was selectively amplified from 1 μl of the infection supernatant in a 50 μl reaction using a primer pair specific to the recombination junction: the first primer (GGTTTCATGTTGAGCAACTGATTCC) targeted the T4^del^ genome outside the recombination site, while the second (TTTGAATTGAAGGAAATTACATGAAACTGTCTAAAGAT) annealed to the integrated plasmid region beyond gp45. This selection strategy ensured that only successfully recombined phage genomes were amplified, effectively excluding non-recombined plasmid and parental T4^del^ genomic background from the final library. These products then served as templates for a second PCR using pool-specific primers to isolate the approximately 200 to 300 bp mutagenized regions, incorporating 5’ overhangs to act as handles for the subsequent attachment of sequencing adapters. In the final stage, unique TruSeq indices were introduced to generate either ∼450bp amplicon or ∼350 bp, which was then sequenced on an Illumina MiSeq platform using MiSeq reagent v2 with 500-cycle kit or 300-cycle kit respectively.

### Sequencing Data Processing and Variant Quantification

Following Illumina sequencing, the raw paired-end reads were processed to reconstruct the full-length amplicons encompassing the mutagenized regions. Read merging was accomplished using the FLASH (Fast Length Adjustment of SHort reads) software utility (Magoč and Salzberg 2011). To ensure high-confidence alignments and minimize assembly artifacts, a stringent minimum overlap parameter of 65 nucleotides was enforced during the merging step.

The frequencies of individual sequence variants within both the unselected baseline and post-selection phage libraries were quantified. To maximize specificity and computational efficiency, read counting was performed via exact string matching. Custom shell scripts leveraging the Unix *grep* command-line utility were utilized to search and iteratively tally the exact occurrences of each target DNA sequence directly from the merged output files (Shell Scripts on GitHub).

To mitigate the impact of random jackpot events (Luria and Delbrück 1943), we calculated fitness values by averaging results across at least three independent experimental replicates. For additional mitigation in helix 𝛼1 libraries, a 20-fold excess of wild-type clamp loader DNA was spiked into the input library. Data visualization and downstream statistical analyses were subsequently performed using custom Python scripts utilizing the pandas, Matplotlib, and Seaborn libraries (Scripts on GitHub). To evaluate fitness distributions across different design templates, individual replicate enrichment values were plotted as frequency histograms. Relying on the standard deviation across replicates provided a robust guide to data reliability; variants exceeding a defined threshold (SD > 1) were filtered out to ensure consistency, and the histograms incorporated a secondary axis displaying the average standard deviation for each fitness bin. To map mutational preferences, variant DNA sequences were translated to amino acids, and substitution frequencies at targeted regions were visualized as annotated heatmaps.

### Protein Frustration Analysis

To identify regions of structural flexibility and energetic stress within the clamp loader, we calculated local energetic frustration using the Protein Frustratometer web server (Parra et al. 2016). We determined the configurational frustration index using the active-state structure of the T4 clamp loader complexed with an ATP analog, primer-template DNA, and the sliding clamp (PDB ID: 8UH7) (Marcus et al. 2024).

### ProteinMPNN designs

Computational designs were generated using ProteinMPNN (Dauparas et al. 2022), templated on two distinct states of the T4 clamp loader: the active conformation (PDB ID: 8UH7), which is bound to an ADP-BeF_3_, primer-template DNA, and the sliding clamp, and the inactive conformation (PDB ID: 8UNH), which is bound to the sliding clamp but lacks ATP. While the active conformation contains four defined ATPase subunits (Subunits B–E), the inactive conformation has only two subunits (Subunits B and C) defined in the asymmetric unit. To handle this discrepancy, designs based on the inactive template were restricted to these two defined subunits, using them as the fixed structural context, whereas active state designs utilized the full four-subunit assembly.

We executed three design strategies across three target regions—Helix 𝛼1, Hinge, and Helix 𝛼7—to evaluate the effect of structural context on design fitness. The first strategy allowed for the redesign of all available subunits in the template simultaneously, while the second and third strategies localized the redesign exclusively to Subunit B or Subunit E, respectively, with all other subunits held fixed. This comprehensive approach yielded a final library of unique sequences for each region: 553 sequences for helix 𝛼1 (228 from the active template and 325 from the inactive), 907 for hinge (452 from active and 455 from inactive), and 908 for helix 𝛼7 (350 from active and 558 from inactive).

For all design trajectories, ProteinMPNN was executed at a sampling temperature of 0.25 (arbitrary unit) to generate approximately 200 distinct sequences per run (Dauparas et al. 2022). The resulting variants were evaluated based on the model’s global and per-residue predicted log-probabilities and the top scoring sequences were selected. Custom Python scripts were utilized to parse the output FASTA files and extract unique sequences for experimental validation.

### Sequence Conservation Analysis and Logo Generation

To assess evolutionary conservation across the AAA+ module of the T4 clamp loader (gp44), a sequence logo was generated using a curated alignment of homologs. The T4 gp44 protein sequence was used as a query for a BLASTp search against the non-redundant protein sequence database to identify approximately 2,000 homologous sequences. To ensure a diverse representation of the protein family and avoid overrepresentation of closely related phage sequences, the dataset was filtered to remove any sequences exhibiting more than 70% sequence identity to the T4 gp44 query which resulted in ∼900 sequences. The resulting filtered sequences were aligned, and a sequence logo was generated using the WebLogo server (Crooks et al. 2004). The height of each amino acid character in the logo represents its relative frequency at that position, scaled by the total conservation (measured in bits) to highlight residues critical for the structural and functional integrity of the clamp loader.

### Contributions

S.N. and J.K. conceptualized the study and drafted the manuscript. S.N. and S.S. designed the high-throughput experiments, which were executed by S.N. Sequence variants were designed using ProteinMPNN by S.N., T.N., and D.K. High-throughput experiments involving ProteinMPNN sequences were designed, performed, and analyzed by S.N. and T.N. J.K. provided supervision and project oversight. S.N., M.O’.D., and J.K. wrote and edited the manuscript. All authors reviewed and approved the final version.

## Acknowledgements

We thank the members of the Kuriyan Lab for their insightful discussions throughout this project. We are grateful to Timothy J. Eisen (University of California, Berkeley) for his critical review of the manuscript and for his valuable guidance on data analysis. We also thank Elizabeth A. Komives (University of California, San Diego) for helpful discussion regarding protein frustration. This work was supported by the National Institutes of Health (NIH) under grant R01-GM144512.

## References

Benkovic SJ, Spiering MM. 2017. Understanding DNA replication by the bacteriophage T4 replisome. J. Biol. Chem. 292:18434–18442.

Bloom JD, Arnold FH, Wilke CO. 2007. Breaking proteins with mutations: threads and thresholds in evolution. Mol. Syst. Biol. 3:76.

Bowman GD, O’Donnell M, Kuriyan J. 2004. Structural analysis of a eukaryotic sliding DNA clamp-clamp loader complex. Nature 429:724–730.

Bryngelson JD, Onuchic JN, Socci ND, Wolynes PG. 1995. Funnels, pathways, and the energy landscape of protein folding: a synthesis. Proteins 21:167–195.

Carver A, Zhang B, Zhang X. 2025. Structures and mechanisms of AAA+ protein complexes in DNA processing. Curr. Opin. Struct. Biol. 92:103056.

Crooks GE, Hon G, Chandonia JM, Brenner SE. 2004.WebLogo: a sequence logo generator. Genome Res. 14:1188–1190.

Dauparas J, Anishchenko I, Bennett N, Bai H, Ragotte RJ, Milles LF, Wicky BIM, Courbet A, de Haas RJ, Bethel N, et al. 2022. Robust deep learning-based protein sequence design using ProteinMPNN. Science 378:49–56.

Deléage G, Roux B. 1987. An algorithm for protein secondary structure prediction based on class prediction. Protein Eng. 1:289–294.

Draghi JA, Parsons TL, Wagner GP, Plotkin JB. 2010. Mutational robustness can facilitate adaptation. Nature 463:353–355.

Engler C, Kandzia R, Marillonnet S. 2008. A one pot, one step, precision cloning method with high throughput capability. PLoS ONE 3:e3647.

Fenton AW, Page BM, Spellman-Kruse A, Hagenbuch B, Swint-Kruse L. 2020. Rheostat positions: A new classification of protein positions relevant to pharmacogenomics. Med. Chem. Res. 29:1133–1146.

Ferreiro DU, Hegler JA, Komives EA, Wolynes PG. 2007. Localizing frustration in native proteins and protein assemblies. Proc Natl Acad Sci USA 104:19819–19824.

Ferreiro DU, Hegler JA, Komives EA, Wolynes PG. 2011. On the role of frustration in the energy landscapes of allosteric proteins. Proc Natl Acad Sci USA 108:3499–3503.

Ferreiro DU, Komives EA, Wolynes PG. 2018. Frustration, function and folding. Curr. Opin. Struct. Biol. 48:68–73.

Fuglestad B, Gasper PM, McCammon JA, Markwick PRL, Komives EA. 2013. Correlated motions and residual frustration in thrombin. J. Phys. Chem. B 117:12857–12863.

Guenther B, Onrust R, Sali A, O’Donnell M, Kuriyan J. 1997. Crystal structure of the delta’ subunit of the clamp-loader complex of E. coli DNA polymerase III. Cell 91:335–345.

Hayden EJ, Ferrada E, Wagner A. 2011. Cryptic genetic variation promotes rapid evolutionary adaptation in an RNA enzyme. Nature 474:92–95.

Hidalgo F, Nocka LM, Shah NH, Gorday K, Latorraca NR, Bandaru P, Templeton S, Lee D, Karandur D, Pelton JG, et al. 2022. A saturation-mutagenesis analysis of the interplay between stability and activation in Ras. eLife 11.

Jayaraman V, Toledo-Patiño S, Noda-García L, Laurino P. 2022. Mechanisms of protein evolution. Protein Sci. 31:e4362.

Jeruzalmi D, O’Donnell M, Kuriyan J. 2001. Crystal structure of the processivity clamp loader gamma (gamma) complex of E. coli DNA polymerase III. Cell 106:429–441.

Jeruzalmi D, O’Donnell M, Kuriyan J. 2002. Clamp loaders and sliding clamps. Curr. Opin. Struct. Biol. 12:217–224.

Kelch BA, Makino DL, O’Donnell M, Kuriyan J. 2011. How a DNA polymerase clamp loader opens a sliding clamp. Science 334:1675–1680.

Kelch BA, Makino DL, O’Donnell M, Kuriyan J. 2012. Clamp loader ATPases and the evolution of DNA replication machinery. BMC Biol. 10:34.

Khan YA, White KI, Brunger AT. 2022. The AAA+ superfamily: a review of the structural and mechanistic principles of these molecular machines. Crit. Rev. Biochem. Mol. Biol. 57:156–187.

Lenzen CU, Steinmann D, Whiteheart SW, Weis WI. 1998. Crystal structure of the hexamerization domain of N-ethylmaleimide-sensitive fusion protein. Cell 94:525–536.

Luria SE, Delbrück M. 1943. Mutations of Bacteria from Virus Sensitivity to Virus Resistance. Genetics 28:491–511.

Magoč T, Salzberg SL. 2011. FLASH: fast length adjustment of short reads to improve genome assemblies. Bioinformatics 27:2957–2963.

Mallam AL, Del Campo M, Gilman B, Sidote DJ, Lambowitz AM. 2012. Structural basis for RNA-duplex recognition and unwinding by the DEAD-box helicase Mss116p. Nature 490:121–125.

Marcus K, Huang Y, Subramanian S, Gee CL, Gorday K, Ghaffari-Kashani S, Luo XR, Zheng L, O’Donnell M, Subramaniam S, et al. 2024. Autoinhibition of a clamp-loader ATPase revealed by deep mutagenesis and cryo-EM. Nat. Struct. Mol. Biol. 31:424–435.

Marzahn MR, Hayner JN, Finkelstein J, O’Donnell M, Bloom LB. 2014. The ATP sites of AAA+ clamp loaders work together as a switch to assemble clamps on DNA. J. Biol. Chem. 289:5537–5548.

Masel J. 2006. Cryptic genetic variation is enriched for potential adaptations. Genetics 172:1985–1991.

Meinhardt S, Manley MW, Parente DJ, Swint-Kruse L. 2013. Rheostats and toggle switches for modulating protein function. PLoS ONE 8:e83502.

Mulkidjanian AY, Makarova KS, Galperin MY, Koonin EV. 2007. Inventing the dynamo machine: the evolution of the F-type and V-type ATPases. Nat. Rev. Microbiol. 5:892–899.

Neuwald AF, Aravind L, Spouge JL, Koonin EV. 1999. AAA+: A class of chaperone-like ATPases associated with the assembly, operation, and disassembly of protein complexes. Genome Res. 9:27–43.

Nolan JM, Petrov V, Bertrand C, Krisch HM, Karam JD. 2006. Genetic diversity among five T4- like bacteriophages. Virol. J. 3:30.

Parra RG, Ferreiro DU. 2025. Frustration, dynamics, and catalysis. Curr. Opin. Struct. Biol. 94:103127.

Parra RG, Schafer NP, Radusky LG, Tsai M-Y, Guzovsky AB, Wolynes PG, Ferreiro DU. 2016. Protein Frustratometer 2: a tool to localize energetic frustration in protein molecules, now with electrostatics. Nucleic Acids Res. 44:W356–60.

Petrov VM, Nolan JM, Bertrand C, Levy D, Desplats C, Krisch HM, Karam JD. 2006. Plasticity of the gene functions for DNA replication in the T4-like phages. J. Mol. Biol. 361:46–68.

Raman AS, White KI, Ranganathan R. 2016. Origins of allostery and evolvability in proteins: A case study. Cell 166:468–480.

Rausch AO, Freiberger MI, Leonetti CO, Luna DM, Radusky LG, Wolynes PG, Ferreiro DU, Parra RG. 2021. FrustratometeR: an R-package to compute local frustration in protein structures, point mutants and MD simulations. Bioinformatics 37:3038–3040.

Rockah-Shmuel L, Tóth-Petróczy Á, Tawfik DS. 2015. Systematic mapping of protein mutational space by prolonged drift reveals the deleterious effects of seemingly neutral mutations. PLoS Comput. Biol. 11:e1004421.

Sosa RP, Florez Ariza AJ, Kim J, Tong A, Kang Z-H, Li A, Kuriyan J, Bustamante C. 2026. The Central Coupler of the AAA+ ATPase ClpXP Controls Intersubunit Communication and Couples the Conversion of Chemical Energy into the Generation of Force. BioRxiv.

Soskine M, Tawfik DS. 2010. Mutational effects and the evolution of new protein functions. Nat. Rev. Genet. 11:572–582.

Story RM, Steitz TA. 1992. Structure of the recA protein-ADP complex. Nature 355:374–376.

Subramanian S, Gorday K, Marcus K, Orellana MR, Ren P, Luo XR, O’Donnell ME, Kuriyan J. 2021. Allosteric communication in DNA polymerase clamp loaders relies on a critical hydrogen-bonded junction. eLife 10.

Subramanian S, Zhang W, Nimkar S, Kamel M, O’Donnell M, Kuriyan J. 2024. Adaptive Capacity of a DNA Polymerase Clamp-loader ATPase Complex. Mol. Biol. Evol. 41.

Swint-Kruse L, Martin TA, Page BM, Wu T, Gerhart PM, Dougherty LL, Tang Q, Parente DJ, Mosier BR, Bantis LE, et al. 2021. Rheostat functional outcomes occur when substitutions are introduced at nonconserved positions that diverge with speciation. Protein Sci. 30:1833–1853.

Treuheit NA, Beach MA, Komives EA. 2011. Thermodynamic compensation upon binding to exosite 1 and the active site of thrombin. Biochemistry 50:4590–4596.

Wagner A. 2005. Robustness, evolvability, and neutrality. FEBS Lett. 579:1772–1778.

Wicky BIM, Milles LF, Courbet A, Ragotte RJ, Dauparas J, Kinfu E, Tipps S, Kibler RD, Baek M, DiMaio F, et al. 2022. Hallucinating symmetric protein assemblies. Science 378:56–61.

Wilmot CM, Thornton JM. 1988. Analysis and prediction of the different types of β-turn in proteins. J. Mol. Biol. 203:221–232.

Yu RC, Hanson PI, Jahn R, Brünger AT. 1998. Structure of the ATP-dependent oligomerization domain of N-ethylmaleimide sensitive factor complexed with ATP. Nat. Struct. Biol. 5:803–811.

Zheng F, Georgescu R, Yao NY, Li H, O’Donnell ME. 2022. Cryo-EM structures reveal that RFC recognizes both the 3’- and 5’-DNA ends to load PCNA onto gaps for DNA repair. eLife 11.

