## Supplementary_figure for "Redesign of energetically frustrated regions rescues function in defective T4 clamp loaders"

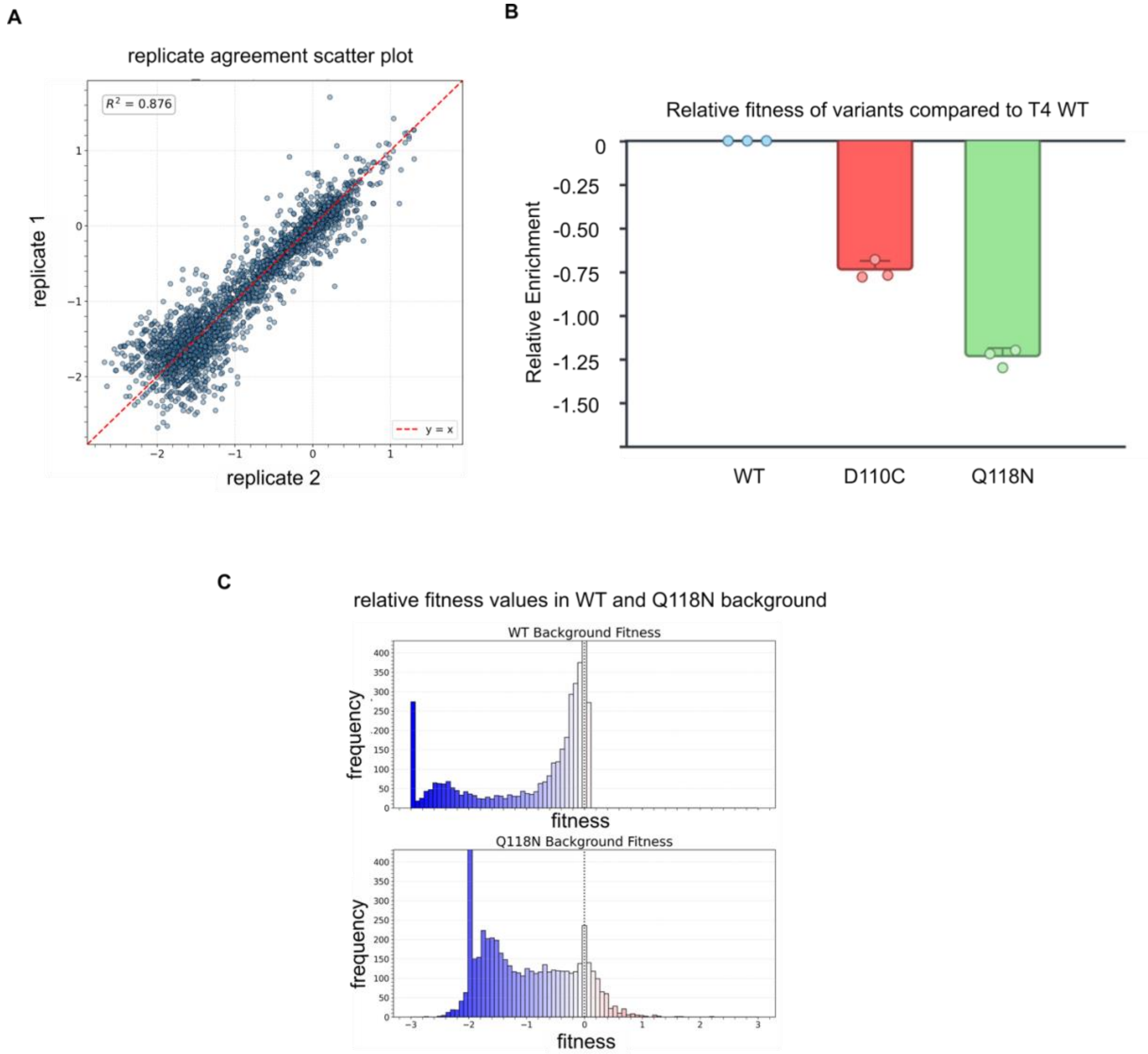


**Figure S1: Fitness distributions of the Q118N saturation mutagenesis screen.**

a) Scatter plot demonstrating high agreement between independent replicates of the Q118N mutant library fitness measurements.

b) Relative fitness of Q118N and D110C variants of T4 clamp loader. D110C and Q118N were found to be approximately 5-fold and 20-fold less fit than the wild-type (WT), respectively.

c) Histograms of mutational fitness effects. Compared to the wild-type clamp loader (top), the Q118N variant (bottom) is globally more sensitive to mutations, exhibiting a larger proportion of deleterious substitutions. Concurrently, the Q118N background reveals a distinct tail of gain-of-function substitutions (fitness > 0, red bars) that act as rescue mutations.

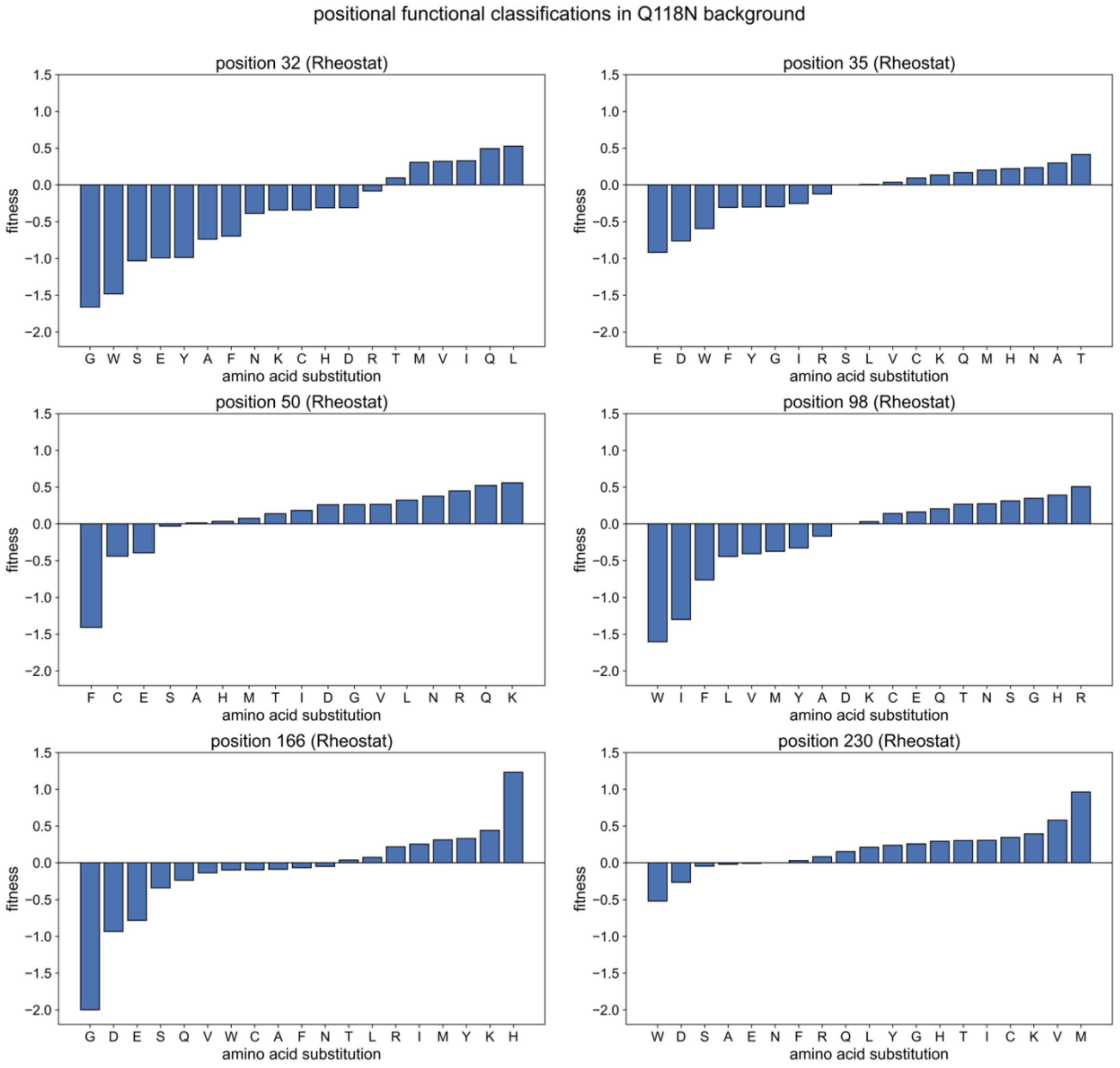


**Figure S2: Rescue hotspots act as conditional rheostats in the Q118N background.** Bar plots detailing the average relative fitness effects of all single amino acid substitutions at representative rescue hotspot positions (residues 32, 35, 50, 98, 166 and 230) within Q118N clamp loader. The horizontal black line at 0 represents the baseline fitness of the Q118N mutant. At each of these positions, different amino acid substitutions yield a wide, continuous spectrum of fitness outcomes, ranging from severe loss-of-function (negative values) to substantial gain-of-function (positive values). This variable, substitution-dependent response defines these sites as "rheostat positions." Because mutations at these same sites are largely neutral in the unperturbed wild-type clamp loader, their rheostatic capacity is conditional. We could not observe any “Neutral” position in Q118N background.

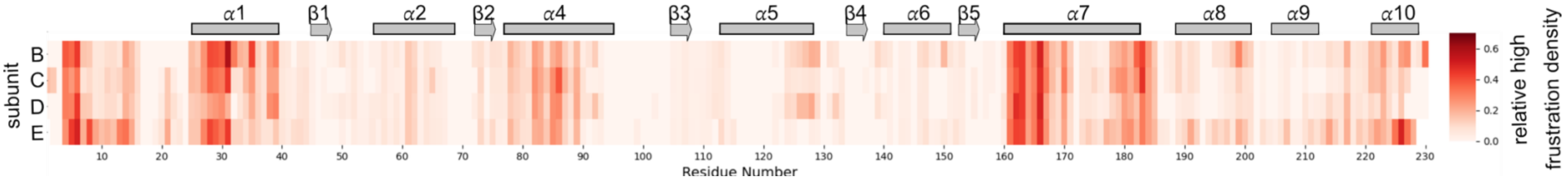


**Figure S3: Configurational frustration in T4 clamp loader.** Configurational frustration for the T4 clamp loader (PDB 8UH7) was calculated using the Frustratometer server. The frustration index profiles for subunits B–E are shown above. Despite the distinct functional roles these subunits perform during the clamp-loading cycle, they exhibit remarkably similar frustration index profiles.

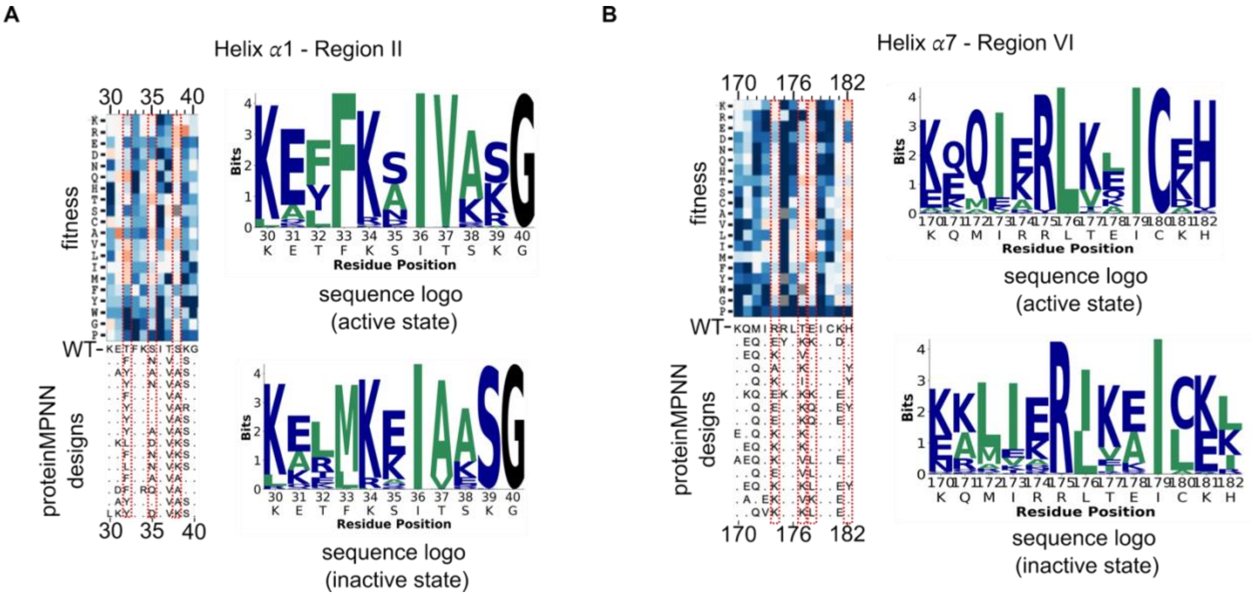


**Figure S4**: **Alignment of ProteinMPNN designs with mutational data for the Q118N mutant for helix $\alpha$1 and helix $\alpha$7.** The heatmap shows fitness data from saturation mutagenesis of the Q118N mutant. Below, an alignment of 15 representative sequences designed using the active structure template is shown. Red dashed boxes highlight rescue hotspots. Sequence logos show the distribution of amino acids in the designed sequences, for active (top) and inactive (bottom) structural templates.

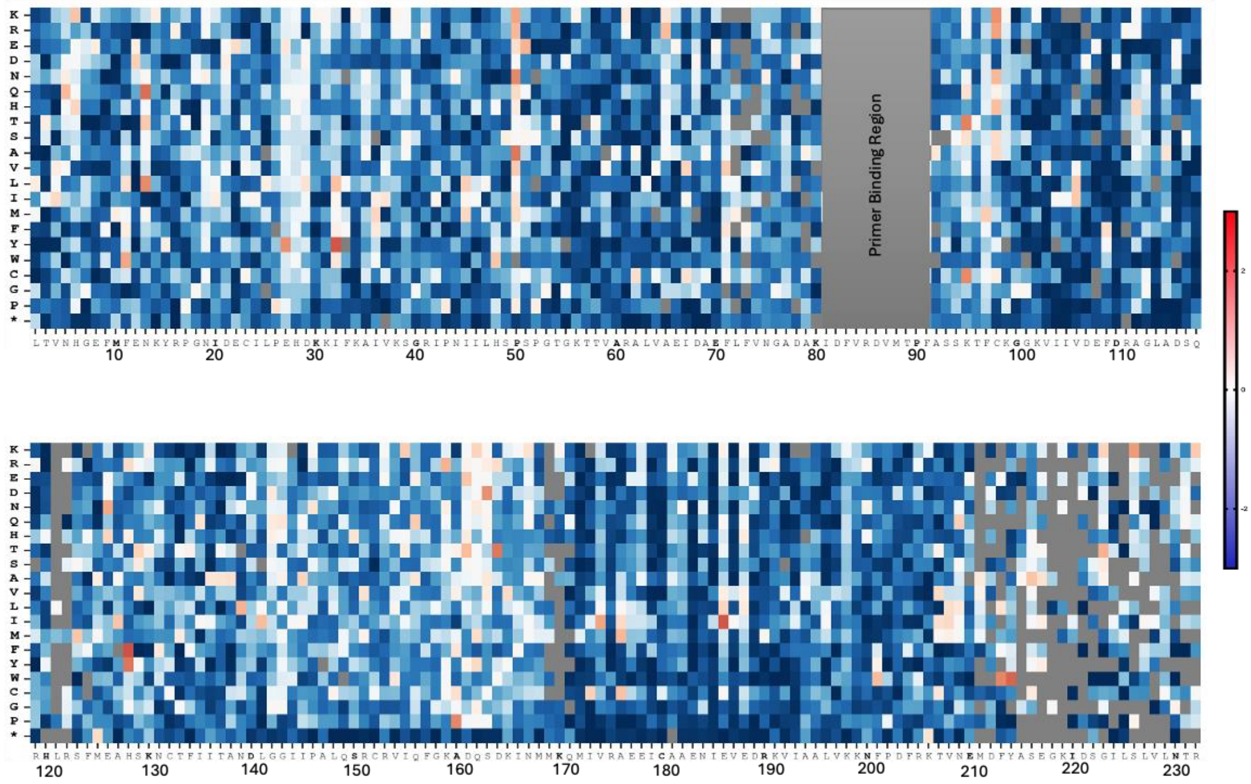


**Figure S5: Deep mutational scanning of the T4/RR2$^{D86H}$ chimeric clamp loader.** The landscape reveals a highly constrained, destabilized complex characterized by widespread deleterious mutational effects and a limited number of rescue hotspots compared to the native T4 mutant variants.

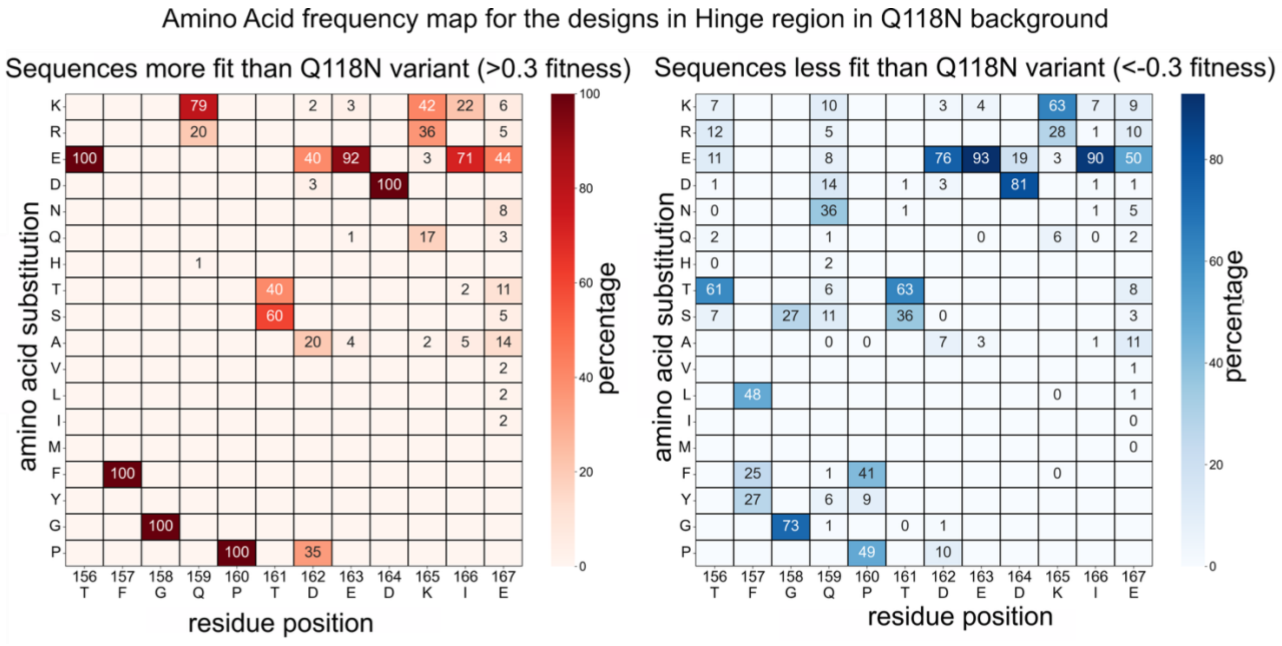


**Figure S6: Sequence determinants of functional rescue in the designed hinge region.**

(Left, red) Frequency matrix for designed sequences that successfully rescue fitness (relative fitness > 0.3). Functional designs converge on highly specific sequence features, most notably the strict conservation of Glu 156 and a strong preference for basic residues (Lys/Arg) at position 159, which are positioned to form a stabilizing ion pair. Additionally, Phe 157 and Pro 160 are strictly conserved (100% frequency) in the working designs. (Right, blue) Frequency matrix for sequences that fail to rescue fitness (relative fitness < -0.3). Non-functional sequences exhibit a disruption of these critical stabilizing features, frequently substituting the crucial E156, F157, and P160 positions with other residues, demonstrating that these specific topological constraints are strictly required to restore the conformational equilibrium.

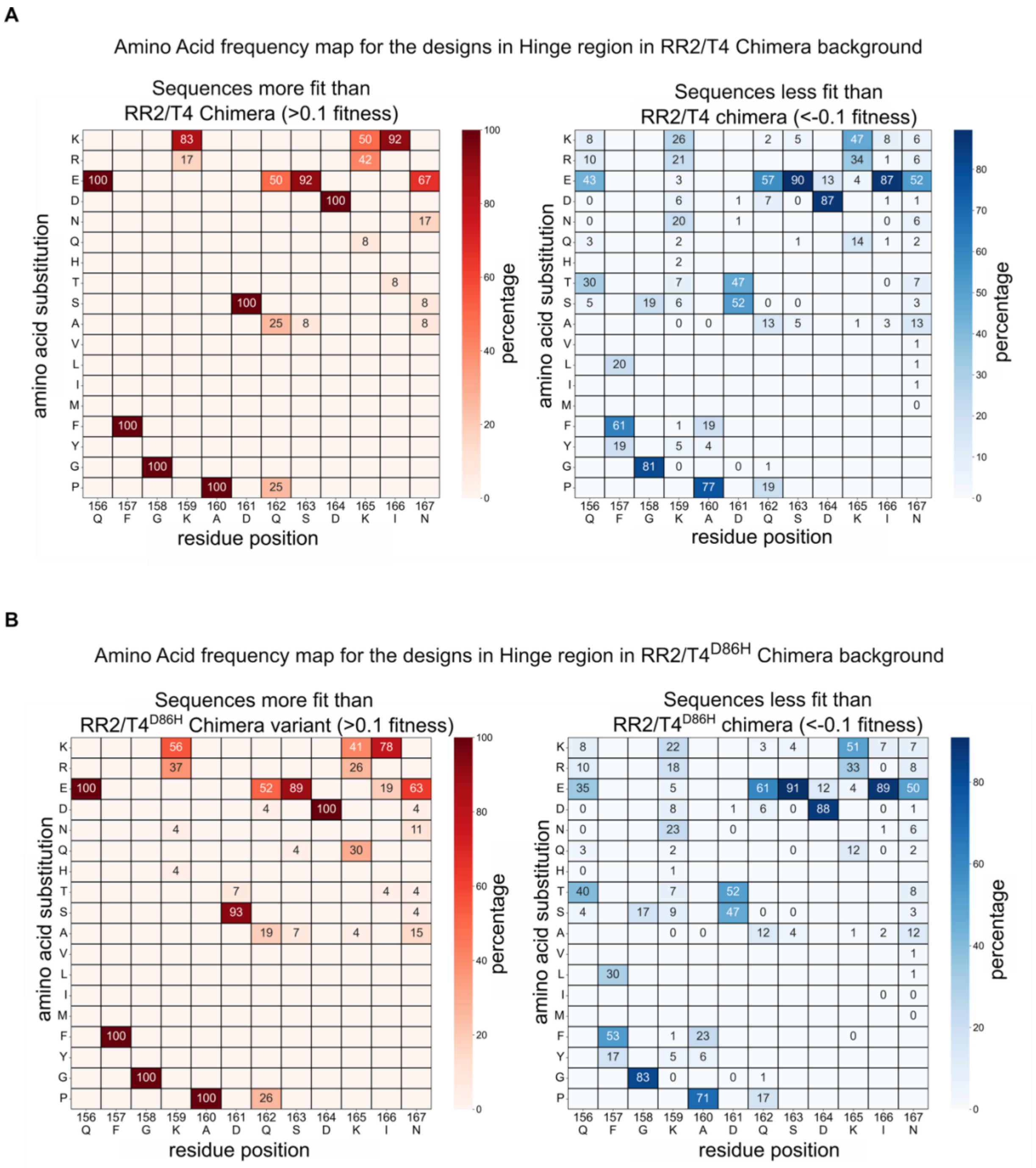


**Figure S7: Sequence determinants of functional rescue are conserved in highly compromised chimeric clamp loaders in the hinge region.**

Amino acid frequency maps for ProteinMPNN-designed hinge sequences (residues 156–167) tested in the T4/RR2 chimera (A) and the T4/RR2$^{D86H}$ chimera (B). (Left, red) Frequency matrices for designed sequences that successfully rescue fitness (relative fitness > 0.1). Consistent with observations in the Q118N background, functional rescue in these severe chimeras is dictated by strict sequence constraints. Successful designs converge on the introduction of a putative ion pair (E156 and K/R159) and the strict conservation of F157, G158, and P160. Notably, the native RR2 AAA+ module possesses an Alanine at position 160; however, functional designs overwhelmingly mutate this position to Proline, mimicking the stabilizing constraint of the wild-type T4 sequence. (Right, blue) Frequency matrices for sequences that fail to rescue fitness (relative fitness < -0.1). Non-functional sequences in both chimeric backgrounds lack these core stabilizing features, exhibiting high mutational variability at the critical 156, 157, 158, and 160 positions.

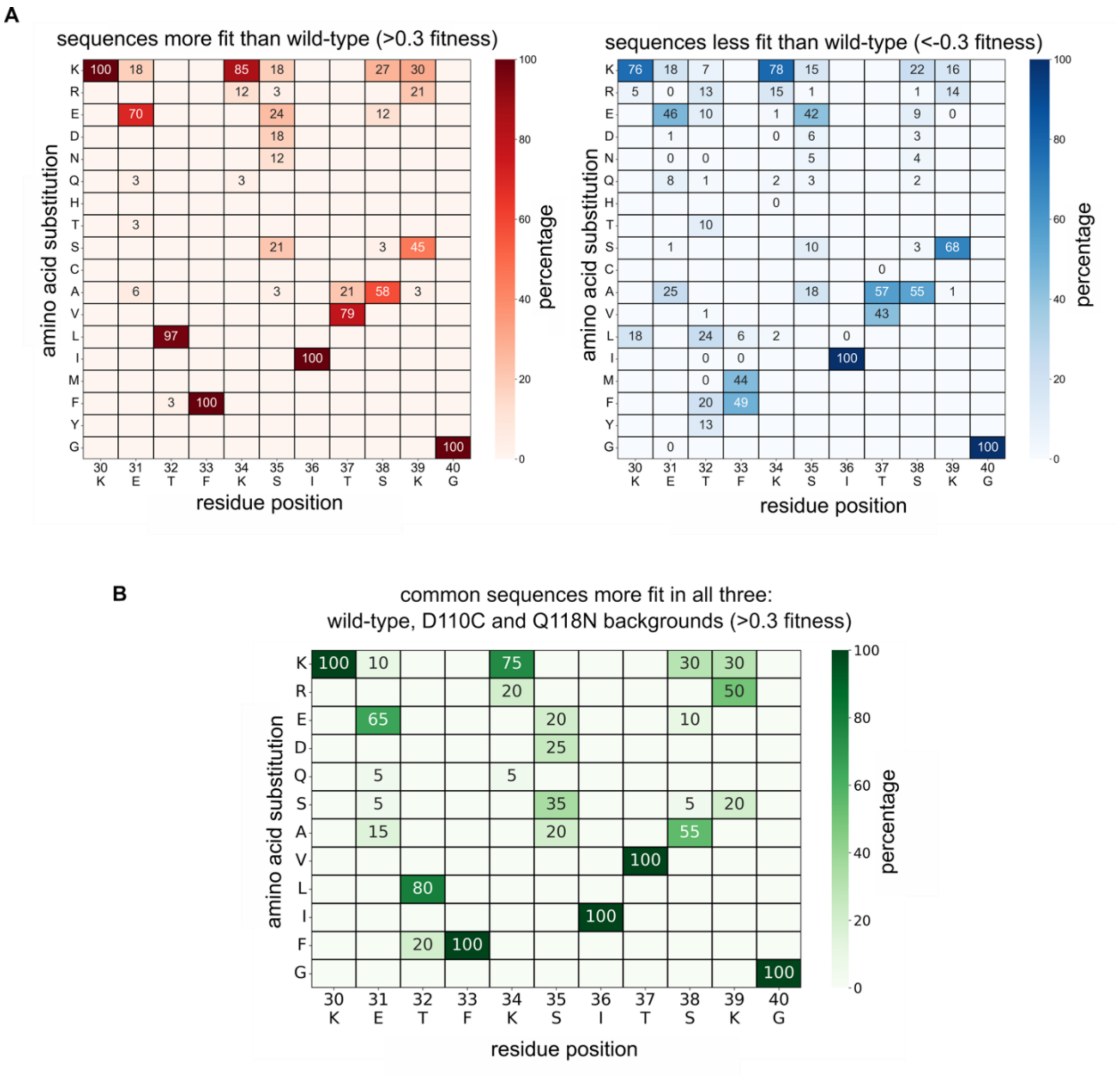


**Figure S8: Position specific amino acid frequency in the helix $\alpha$1 design sequences.**

a) Frequency matrix for designed sequences demonstrating a gain of fitness relative to the wild-type clamp loader (left). High-fitness designs show strong preferences for specific stabilizing substitutions, notably T32L and T37V, while strictly conserving native residues such as K30, F33, I36, and G40. Frequency matrix for designed sequences exhibiting a loss of fitness relative to wild-type (left). Deleterious sequences display distinct, unfavorable mutational profiles (e.g., introducing T37A or K39S) that disrupt the optimal helical packing. b) Frequency matrix isolating "universal" high-fitness designs: a consensus profile of sequences that simultaneously achieve a fitness > 0.3 across the wild-type, Q118N, and D110C backgrounds. These robustly stabilizing designs converge on a highly specific sequence motif, rigidly incorporating the T32L and T37V substitutions alongside the strictly conserved native topological constraints.
